# Flexibility of Short-chain dehydrogenase is interconnected to its promiscuity for the reduction of multiple ketone intermediates

**DOI:** 10.1101/2021.07.05.449867

**Authors:** Anirudh P Shanbhag, Sreenath Rajagopal, Arindam Ghatak, Nainesh Katagihallimath, Ramswamy S., Santanu Datta

**Author notes:** **Corresponding Author:** Dr. Santanu Datta, Ph.D.

## Abstract

Short-chain dehydrogenases/reductases (SDRs) are a convenient class of enzymes used to synthesize enantiopure alcohols. Several studies describe native or engineered SDRs for converting substrates of interest using cost and time-intensive high-throughput approaches. The classification of SDRs is based on chain length and cofactor binding site. Of these, the shorter ‘Classical’ and the longer ‘Extended’ enzymes participate in ketoreduction. However, comparative analysis of various modelled SDRs reveals a length independent conserved N-terminal Rossmann fold and a variable C-terminus region for both types. The general hypothesis is that the latter domain influences the enzyme’s flexibility that may affect the observed promiscuity of the enzyme. We have used a machine learning algorithm on this flexible domain to build a rationale to screen promiscuous candidates. We have built a data set consisting of physicochemical properties derived from the amino-acid composition of enzymes to select closely associated promiscuous mesophilic enzymes. The resulting in vitro studies on pro-pharmaceutical substrates illustrate a direct correlation between the C-terminal lid-loop structure, enzyme melting temperature and the turnover number. We present a walkthrough for exploring promiscuous SDRs for catalyzing enantiopure alcohols of industrial importance.

## Introduction

Short-chain dehydrogenases (SDRs) form a promiscuous class of enzymes that utilize NAD(P)H as a cofactor for catalyzing a wide array of prochiral ketones into enantiopure alcohols. These enzymes have been studied by techniques employing site-directed mutagenesis, site saturation mutagenesis, substrate channeling, and gene shuffling. (1)(2)(3). SDRs encompass functions like detoxification, hormone metabolism, secondary metabolite and biopolymers production (4)(5)(6)(7). Such broad substrate promiscuity is often explored to prospect an enzyme with superior kinetic parameters for converting substrates of industrial importance. Many studies have reported engineered enzymes demonstrating better stability, stereoselectivity and turnover characteristics. However, these upgrades often focus on a single substrate of interest rather than a generic improvement with other substrates of similar markush structures (8)(9). Furthermore, substrate catalysis and turnover rates are trade-offs unless the enzymes are specifically engineered (10). Therefore, it is essential to analyse various enzymes to obtain a candidate to convert multiple substrates at a higher turnover.

Structurally, SDRs have a ubiquitous cofactor binding Rossmann fold which is similar to other dehydrogenases such as Medium-chain dehydrogenases, Iron-containing alcohol and Aldehyde dehydrogenases, signifying convergent evolution (11)(12)(13). They are classified into five types, namely, Classical, Extended, Intermediate, Complex and Divergent. Most of them fall under the first two classes and contain the active site SxnYxxxK. However, ‘Classical’ SDRs are ∼250 residues long with TGxGx3G motif as a cofactor binding region. In contrast, the ‘Extended’ counterparts have an Alanine instead of the third Glycine and are ∼350 residues long. Interestingly, another study has analysed thousands of members showing a length independent and extensively interconnected network of similar proteins (14). Such a superfamily with closely related members with natural substrate promiscuity is ideal for understanding multiple substrate catalysis.

SDRs being a large superfamily of proteins (with a monotonically increasing number of members every year), screening for a promiscuous candidate could be technically overwhelming. However, machine learning algorithms have helped decipher patterns among complex data sets and protein data (15)(16). Previously, SDRs have been classified using Hidden Markov Models (HMMs), which are prominently used in protein sequence classification in SCOP, CATH and Uniprot databases (17)(18). The algorithm has powerful statistical functions and can handle variable lengths of protein sequences inputs. However, it has many unstructured parameters and cannot capture higher-order correlations among amino acids in a protein. Also, HMM has to be reasonably constrained and iterative to represent a small fraction of space over a vast number of sequences. Hence, it is optimal in identifying active sites, secondary structure similarities (used in multiple sequence alignment) and native substrates based on existing data.

Additionally, HMM is often unable in sorting proteins based on other overarching factors such as physicochemical properties, taxonomical origins and other non-canonical classifiers. Therefore, instead of engineering a protein for substrate-specific conversion, we have focused on finding a promiscuous SDR which can convert multiple substrates of interest. We wish to classify SDRs using machine learning algorithms to generate a ‘signature’ of multi-substrate catalysis. Unlike traditional prospecting that uses phylogenetic classification to ascertain the closest homolog to a reported promiscuous enzyme, we classify the proteins based on their respective phylum. This strategy eliminates outliers and gives a homogenous dataset for further clustering to ascertain promiscuous candidates based on their proximity to reported ones. Here, we present a rationale and a ‘Proof of Concept’ for deriving an enzyme that can catalyze different active pharmaceutical intermediates.

## Results

### (i) Modelling and architecture of SDRs

Modelling different sized Short-chain dehydrogenases can help in understanding the architecture of the proteins. There is a soft hypothesis that the lid-loop structure, a helix-loop helix-motif, plays an essential role in the enzyme’s flexibility. It is most often the α7 or α6 helix in SDRs. It is a well-characterized structural entity and was first reported in Thiohydorxy naphthalene reductase from *Magnaporthae grisiae*. (19)(20). The dynamic closing of the lid loop upon cofactor and substrate binding restricts substrate access to the active site to a direction that is compatible with the observed stereochemical outcome of ketoreduction (21). We modelled SDRs of varying lengths to determine the influence of the C-terminus on the overall architecture of the protein. Classical and Extended SDRs exist as dimers (or tetramers) and monomers, respectively. However, the models illustrate a different narrative. The N-terminal Rossmann fold is structurally conserved, whereas; the C-terminal forms a lid-loop structure and consists of residues that participate in oligomerization. Some of the modelling data from the ‘longer’ Classical SDRs from other organisms showed a change in loop architecture and the oligomerization of the enzymes (Figure 2).

**Figure 1:**
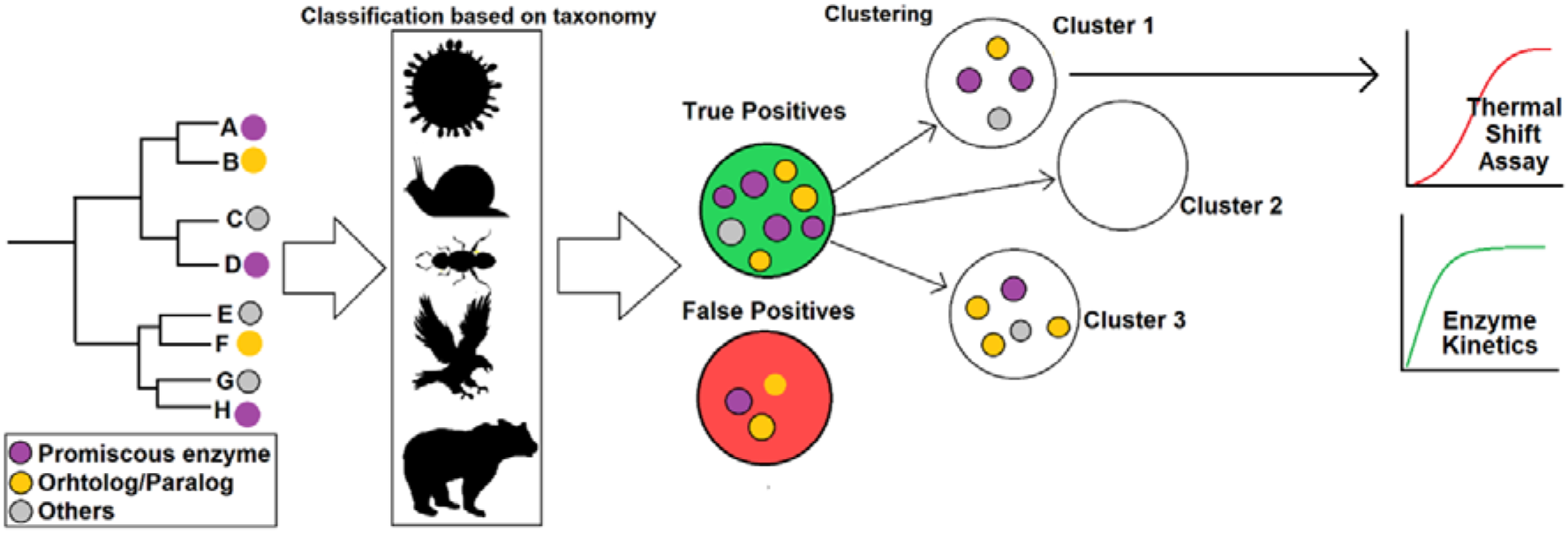
Schematic explaining the prospecting of enzymes using machine learning for clustering candidates with broad substrate activity. Standard phylogenetic classification helps in finding orthologs/paralogs but may not classify enzymes for substrate activities. Due to the similarity in enzymes the short-chain dehydrogenases are classified using supervised learning algorithms to denote true and false positives. Subsequently, they are clustered into enzymes that have a higher probability of being promiscuous. Unknown members closer to the reported promiscuous candidates are purified for in vitro analysis.

**Figure 2:**
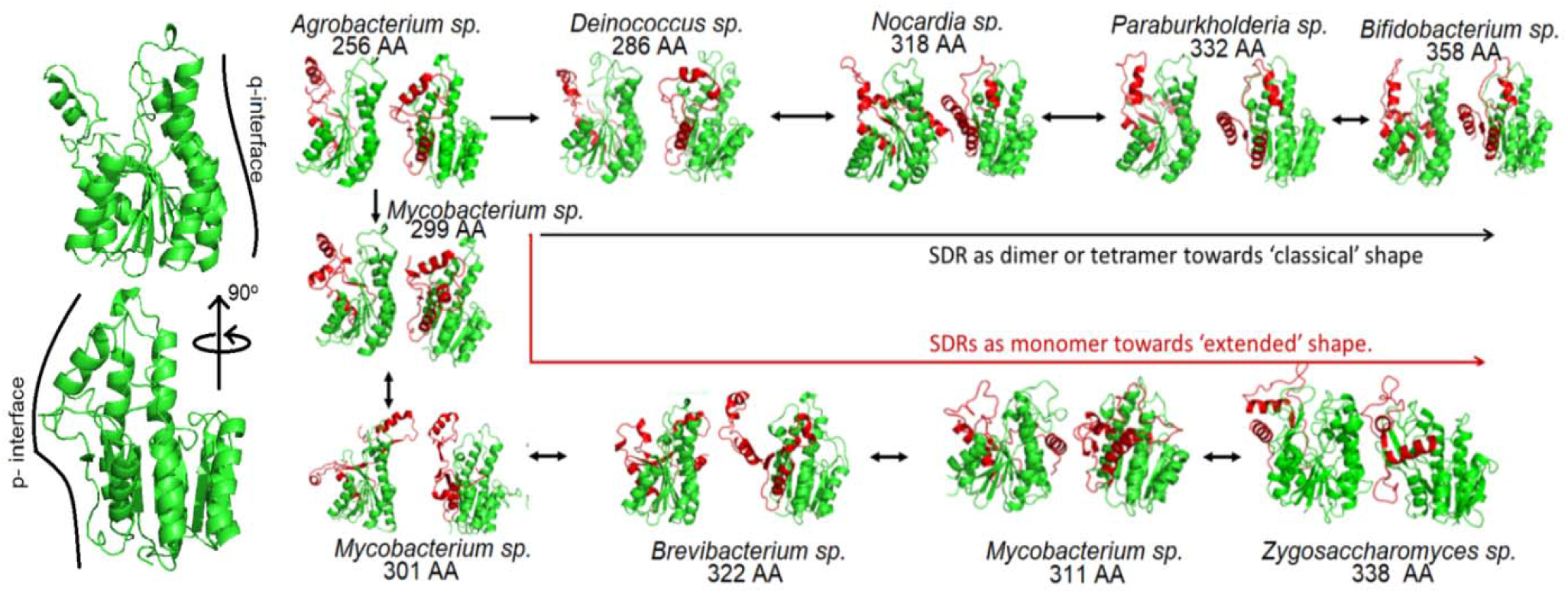
The changes in the lid-loop of the Short-chain dehydrogenases are invariably based on the amino acid composition of the loop affecting the overall architecture and oligomerization of the enzyme. They can exist as Dimers (subsequently Tetramers) in Classical type enzymes and as Monomers in Extended type enzymes

Upon modelling, two distinct sets of SDRs were observed, based on the accessibility of α-helices. Invariably the amino acid composition of the protein is a determines this characteristic. In SDRs which are monomers, the α-helices are not accessible, thus preventing oligomerization. Furthermore, the C-terminal part of the protein participates in changing its shape. Figure 2 demonstrates this change from *Mycobacterium* sp. 301 AA to *Mycobacterium* sp. 311 AA, where the C-terminal loop goes around the P interface (Tetramerization interface) towards the Q interface (Dimer interface), thereby changing the exposure of the α-helix residues imperative for oligomerization. The significance of the orientation of both interfaces was demonstrated by Biswas *et al*., where swapping of the Q-interface domains between the naturally tetrameric *Staphylococcus* FabG and dimeric *Escherichia coli* FabG led to a tetrameric chimeric variant of *E. coli* FabG (22). Another important fact is that the active site is more accessible due to a floppy lid-loop structure (red).

Further, upon comparison with an extended SDR, it is seen that both accessibilities of the active site of the non-accessibility of the Q-interface are evident (Figure 2 *Z. rouxii* 338 AA). In classical SDRs, the α4 and α5 helices are unrestricted and available for oligomerization. However, in extended SDRs, the helices are hindered and therefore, oligomerization fails to occur in these enzymes (23). The change in the cofactor binding site of the Extended SDRs is not a product of the protein’s architecture; instead, it is the composition of the Amino acids. The active site is nearly the same among all the classical SDRs, as seen in Supplementary figure S2.

### (ii) The catalytic efficiency of an essential SDR and its homologs from *E. coli*

FabG is one of the essential SDRs along with the FabI gene in E. coli. Most essential enzymes are specific and have low promiscuity, and therefore, we used the enzyme as a control. UcpA and IdnO are the closest non-essential counterparts of FabG in the organism. They were selected to discern the trend towards multi-substrate catalysis and delineate the relationship between gene essentiality, flexibility, and catalytic promiscuity. These enzymes are similar to each other (Figure 3B). Further, the ketoreductase ZRK was used as a positive control for ascertaining the spectrum of substrate promiscuity.

**Figure 3:**
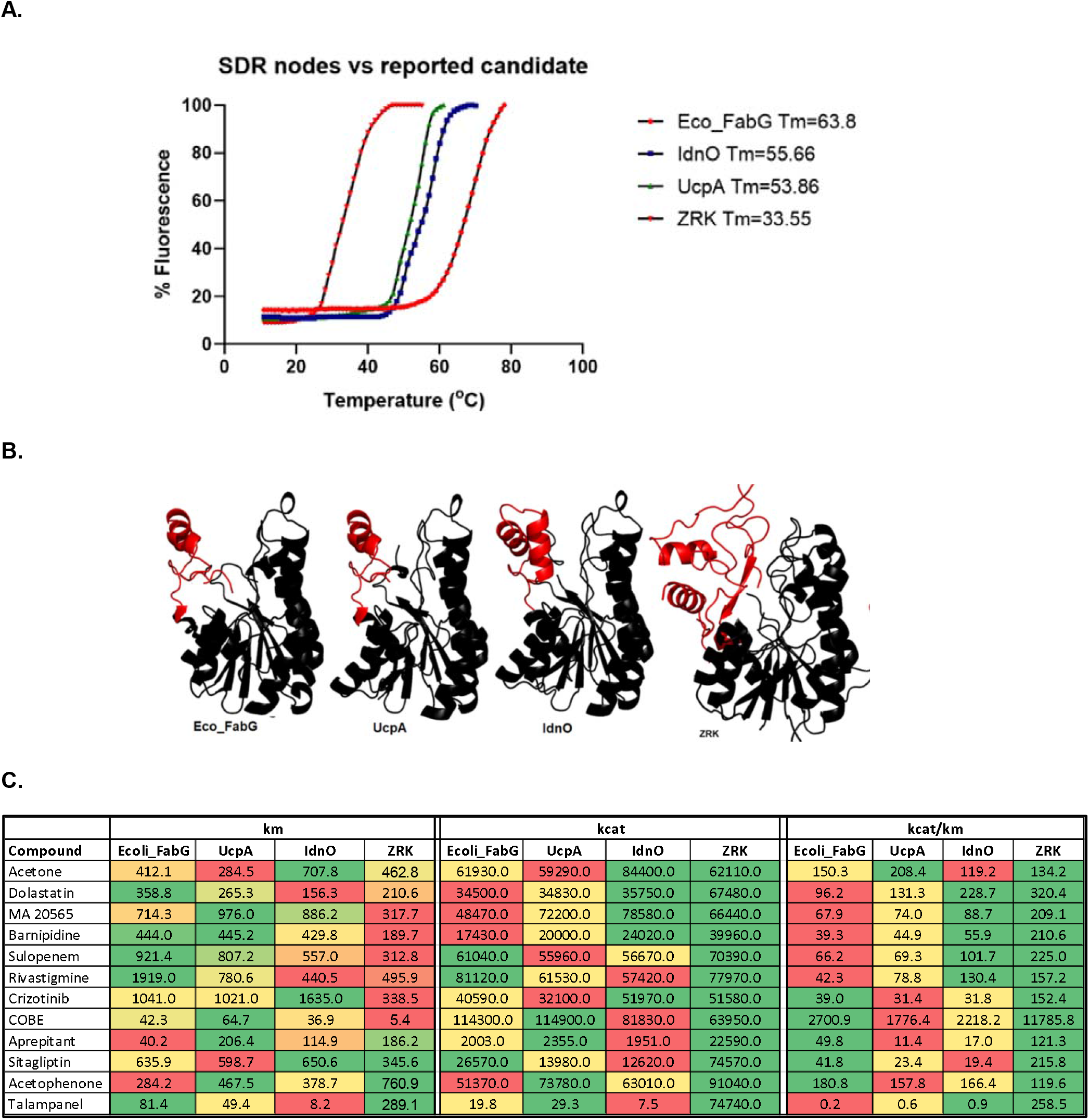
Repurposing of ketoreductases for production of chiral alcohols. A. Differential scanning fluorimetry of short chain dehydrogenases. B. Lid-loop structure (red) of chosen SDRs. C. Heat map of the catalytic efficiency of chosen SDRs Units: km=uM kcat=uM/min.

Differential scanning fluorimetry (Thermal shift assay or DSF) was performed to test the enzymes’ rigidity, where the flexibility of the enzymes is inversely proportional to the melting temperature of the proteins (24). Many reports on enzymes belonging to the same family have demonstrated protein flexibility as a characteristic dependent on the optimum functional temperature. In psychrophiles, the cold-adapted proteins Uracil DNA glycosylases and Endonucleases I are more flexible than their mesophilic counterparts (25)(26). Similarly, thermophilic proteins such as malate dehydrogenases were studied protein rigidity using both differential scanning fluorimetry and Molecular Dynamics (MD) simulations (27)(28).

The dynamicity of the SDR models were checked using various methods. The B-factor was measured on AlphaFold predicted models in ChimeraX tool (Supplementary Figure S4A). Further, the deep-learning server Medusa showed an increase in number of flexible residues in proteins with larger lid-loop structure (Supplementary Figure S4B). We also cross validated this using both the primary sequence and AlphaFold models using PredyFlexy and FlexServ (Supplementary Figure S4 C and D). Thus, it is not overly optimistic to assume that protein flexibility can be further augmented by Differential scanning fluorimetry data. During a DSF assay, the buried hydrophobic residues bind to SYPRO orange as the protein unfolds due to increasing temperature, producing a melt curve. The slope of the melt curve and the midpoint of transition (Tm) indicate the enzyme’s overall rigidity/flexibility (Figure 3A). The models depict a relationship between the lengths of the lid-loop and the proteins’ melting temperature (Figure 3B). The Eco_FabG is relatively rigid and has the highest melting temperature (63.8 °C), while UcpA and IdnO has a lower Tm (54-56 °C), and ZRK has the lowest (33.55 °C) melting temperature (Figure 3A). Furthermore, ZRK shows the highest catalytic efficiency, followed by IdnO and UcpA. Therefore, Eco_FabG, which has the minimum catalytic efficacy, establishes an inverse correlation between rigidity as a characteristic and multi-substrate catalysis (Figure 3C).

### (iii) Machine learning

#### a. Dataset preparation

The dataset was prepared by downloading SDR sequences from UniProt, CATH and SCOP databases. They were then filtered using the Batch CDD (Cluster Domain Database) to identify the family and superfamily, respectively. Furthermore, the taxonomic origin for every protein sequence (at the phylum level) was manually curated while eliminating entries with unknown origins. The features from the ProtScale and ProtParam (ExPasy server) were used to build the dataset. A total of 81,014 curated SDRs were finally deployed for classification.

#### b. Feature selection

In data processing, intelligent feature selection is an essential step for filtering out redundant or low-affecting factors. This step increases the efficiency of sorting through supervised or unsupervised learning. A Chi-square test was performed, and a correlation matrix was produced (Figure 4). Theoretically, a very low or a high number of features often leads to wrong classification or overfitting. In this study, the correlation matrix helped determine 20 features for classifying SDRs; namely, Molecular Wt., GRAVY, the amino acids A, C, D, E, F, G, I, K, L, N, Q, R, S, V, Y, Aliphatic Index, % Negatively charged amino acids, % of Positively charged amino acids were chosen as features for classification.

**Figure 4:**
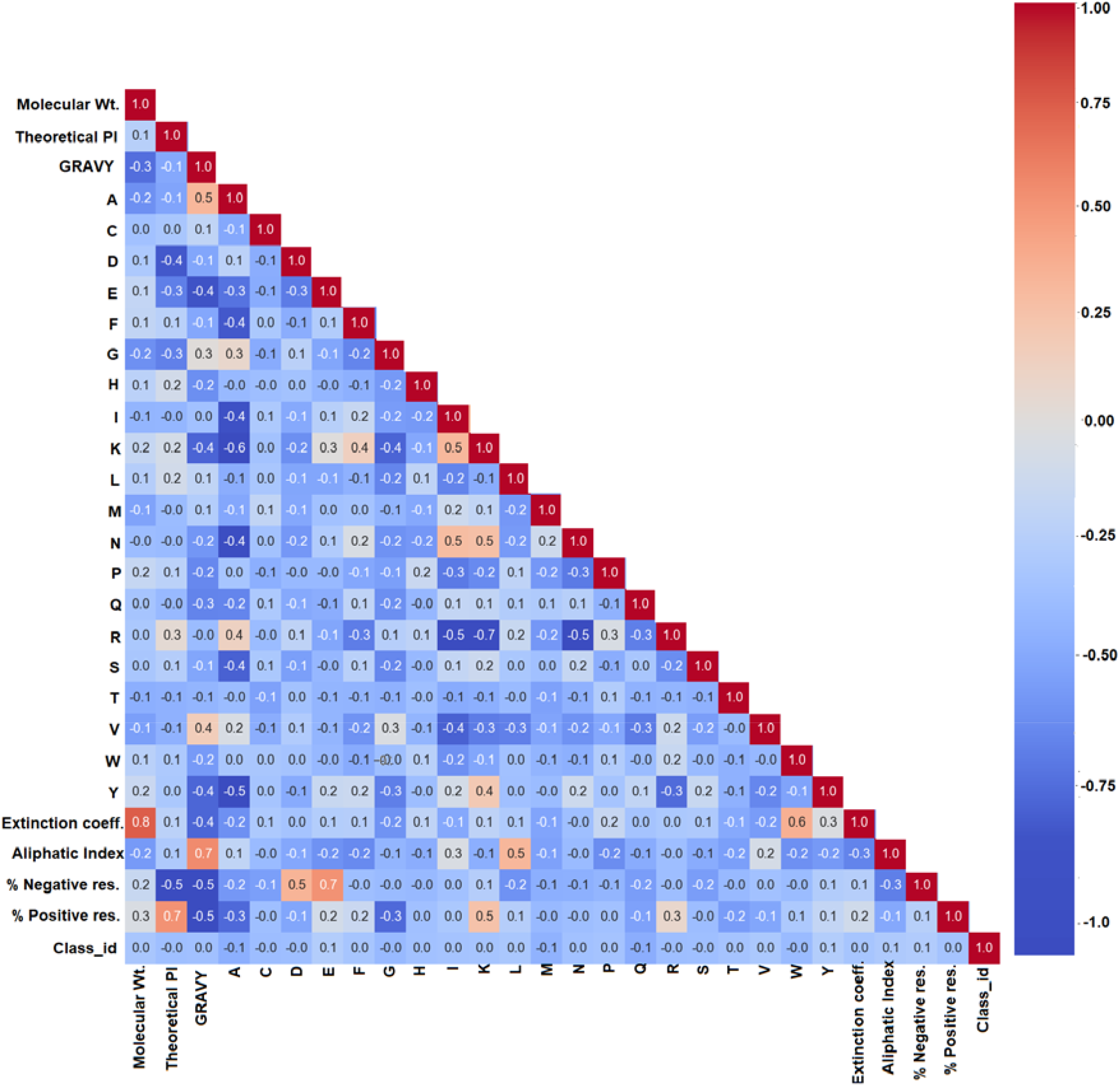
Feature selection of the physicochemical properties using correlation matrix for classifying SDRs. Positive integers depict a positive correlation and negative integers represent vice versa. Top 20 physicochemical properties having correlation values farther from zero were chosen for classification.

#### c. Classification of enzymes

A refined dataset exemplifies the accuracy and performance of the algorithms in question. To this end, the proteins must be refined by considering the gradual change influenced by speciation. Genes (and proteins by extension) are known to evolve from gene duplication, gene transfer and mutation events. Compared to these two occurrences, gene insertions are rare (29)(30)(31). Therefore, classifying sequences based on their phylum of origin can help obtain candidates that have undergone a gradual change in the line of evolution and speciation. SDRs have many members due to gene duplication and translocation events throughout various phyla across geological time. Hence, considering enzymes that have existed in the line of evolution would enable in eliminating strong outliers (which have arisen from gene insertion or multiple gene truncation events) from the dataset. Additionally, most of these events are across closely related species in a phylum, subphylum, and class. Thus, classification was done to avoid extraordinary circumstances in which these genes which might have undergone major deletions or insertions. This process refines the dataset for the subsequent clustering analysis.

Machine learning algorithms such as k-Nearest Neighbour, Extra tree, Random Forest, and Support Vector machines were used for classifying enzymes to their respective derivative organism’s phyla based on the physicochemical parameters. This process helped retain only true positives using the confusion matrix (Example shown in Figure 5A and complete data is depicted in Supplementary figures S7-S14). We observed that True positives were (marginally) better classified using Support vector machines (SVM) than other data-mining algorithms. Each machine learning algorithm underwent 1500 iterations, beyond which the accuracy could not be improved to greater than 80% for SDRs (Figure 5B).

**Figure 5:**
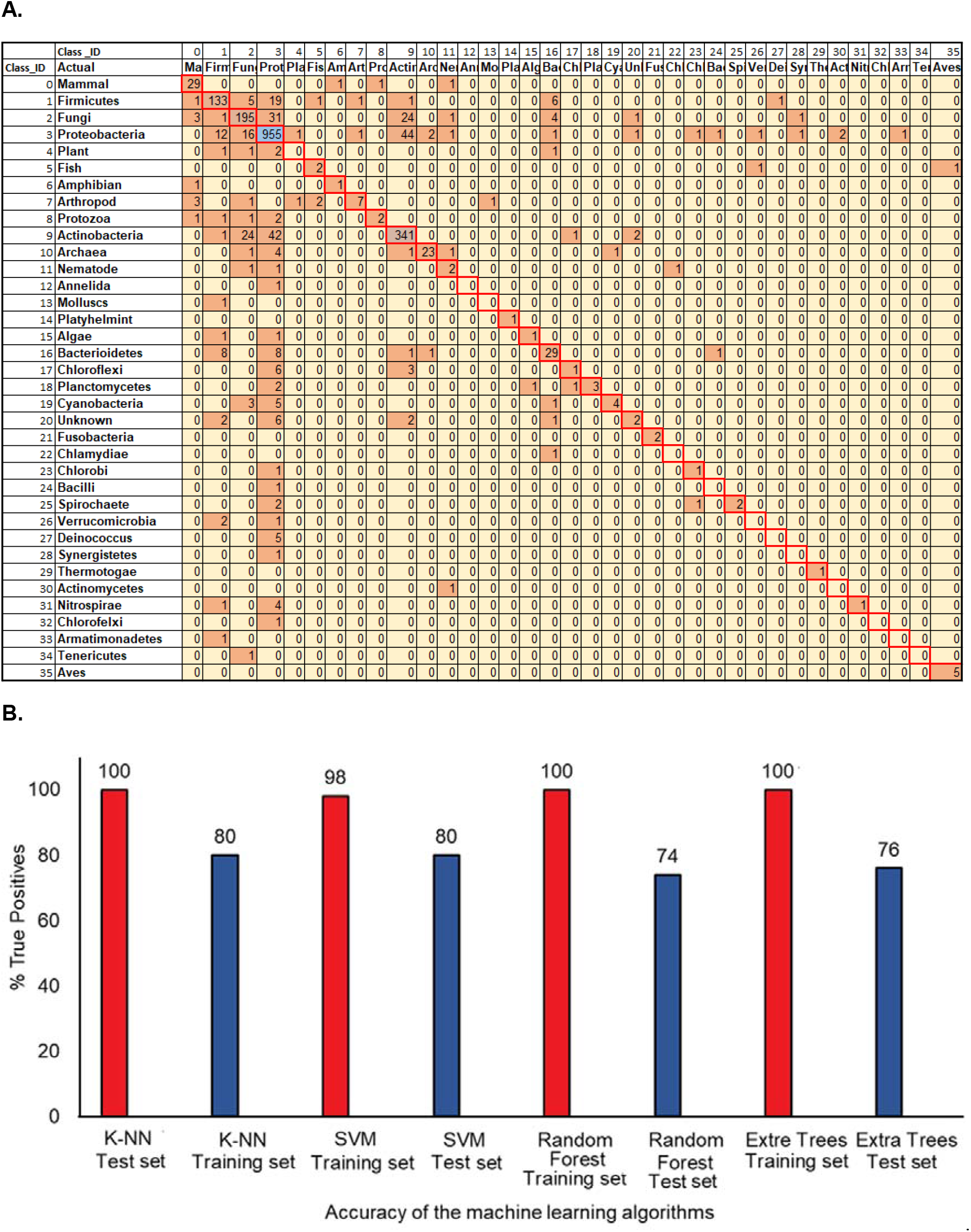
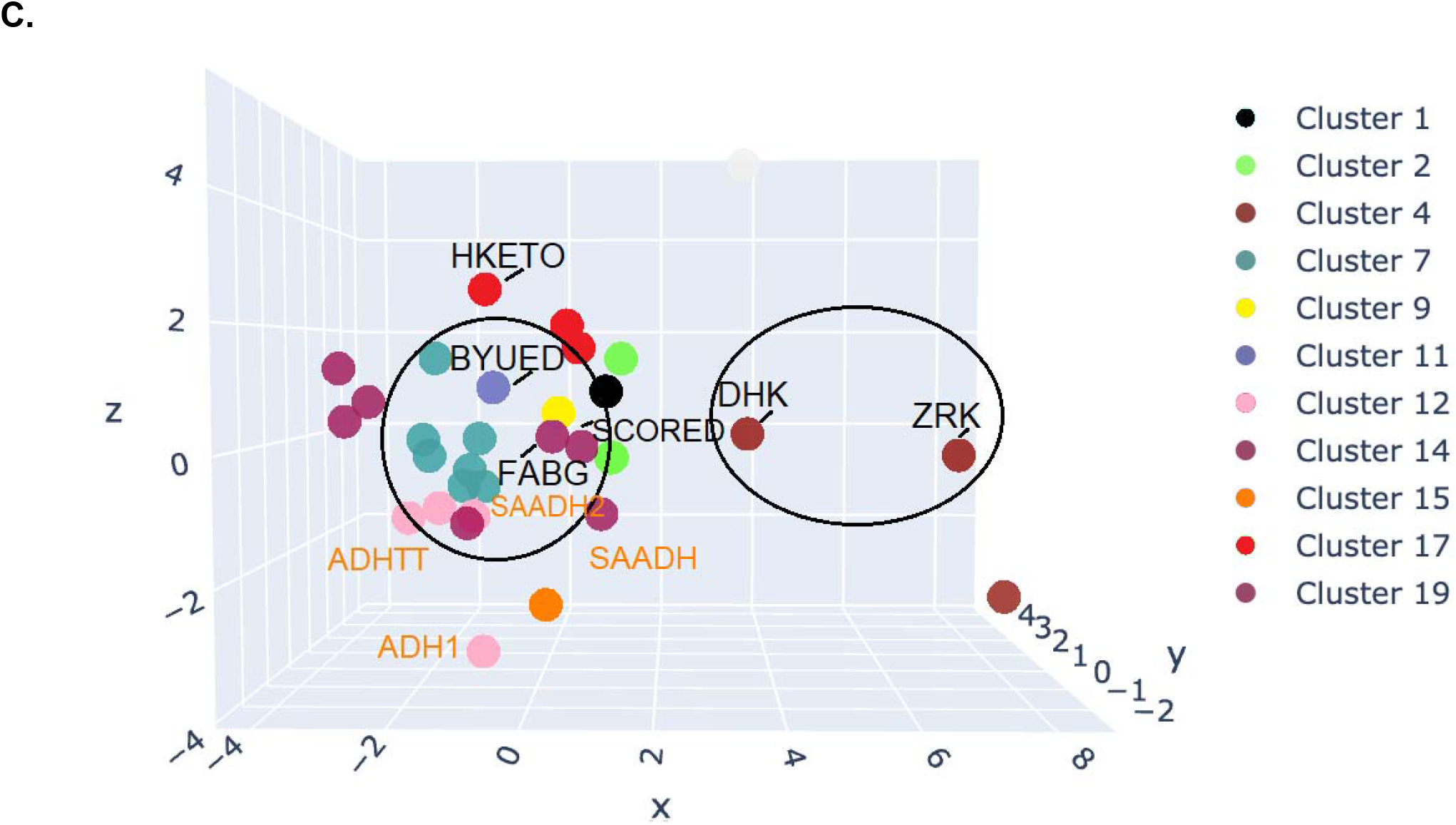
A. An example of a confusion matrix portraying true positives (reverse diagonal marked in red) obtained by SVM for SDRs. B: Graph depicting the accuracy of Support vector machines (SVM), Extra trees, Random Forest, and k-Nearest Neighbour method in classifying SDRs based on physicochemical parameters. C. Principal component analysis (PCA) of SDRs where the chosen members for in vitro analysis are encircled, and the thermophilic enzymes are depicted in orange

#### d. Clustering of enzymes

The number of clusters was determined using the Elbow (or knee of a curve) method, a heuristic approach for determining the number of optimum clusters for any given dataset. K-Means clustering was used for finding enzymes that lie close to the reported promiscuous enzymes. Since k-means clustering uses a mean between three data points, i.e., the cluster’s centroid defined as epsilon value (the sum of squares of the distance between data points and the respective centroid of the cluster to which the data point belongs) was measured using the squares of the distances between them. The elbow method uses a two-dimensional plot between the epsilon value and the number of iterations to determine the number of clusters (Supplementary figure S5).

The clustering results show that enzymes group together based on their catalytic potential (promiscuity) and melting temperature. Thermophilic enzymes belonging to clusters 7, 12 and 19 are close to each other. We used principal component analysis (PCA) for graphically depicting these results (Figure 5C). Similarly, clusters 4, 7, 11, 12, 19 have promiscuous enzymes grouped in the neighbourhood. Except for clusters 12 and19, most of the enzymes from the remaining clusters contain promiscuous candidates. Potential unknown enzymes clustered with reported promiscuous SDRs were chosen for in vitro analysis (Table 1). ByueD (*Bacillus subtilis* ketoreductase) and DHK (*Debaryomyces hansenii* ketoreductase) were selected for further analysis. As a negative control, another unverified enzyme Hketo (*Hansenula polymorpha* ketoreductase) was chosen from cluster 17 (a low promiscuous cluster in Table 1).

**Table 1:**
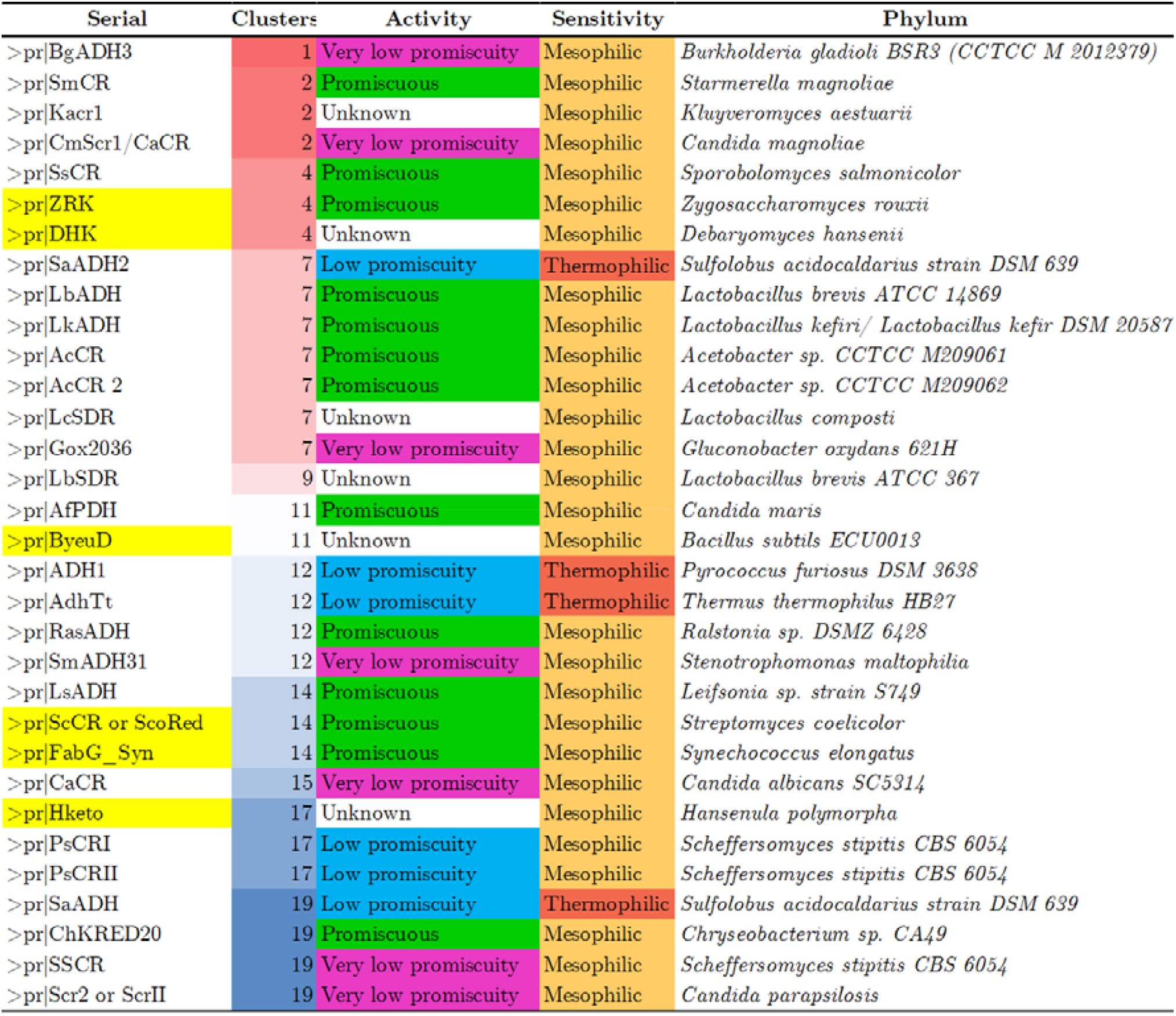
Clustering of the reported 23 SDRs from the total 81,014 members. Characteristics such as promiscuity and sensitivity are present in the same clusters. Some of the unknown enzymes (highlighted in yellow) lying in clusters having promiscuous and non-promiscuous members were chosen for in vitro analysis.

### (iv) In vitro analysis

For In vitro analysis, enzymes were chosen from the clusters based on the different architecture. As aforementioned, SDRs chosen for ketoreduction are predominantly from classical and ‘extended’ types. Therefore, from the cluster, three known and three unknown enzymes were chosen for in vitro analysis. Classical SDRs are known for producing Active Pharmaceutical Intermediates (API), namely, *Synechococcus sp*. PCC 7094 FabG, ScoRed (having a classical SDR structure similar to Eco_FabG) were selected, and ZRK were chosen from the cluster. Additionally, candidates not reported for the production of Active Pharmaceutical Intermediates such as ByueD (*Bacillus subtilis)*, Hketo *(Hansenula polymorpha)* and DHK (*Debaryomyces hansenii*) were chosen based on the clusters predicted (Figure 6B).

**Figure 6:**
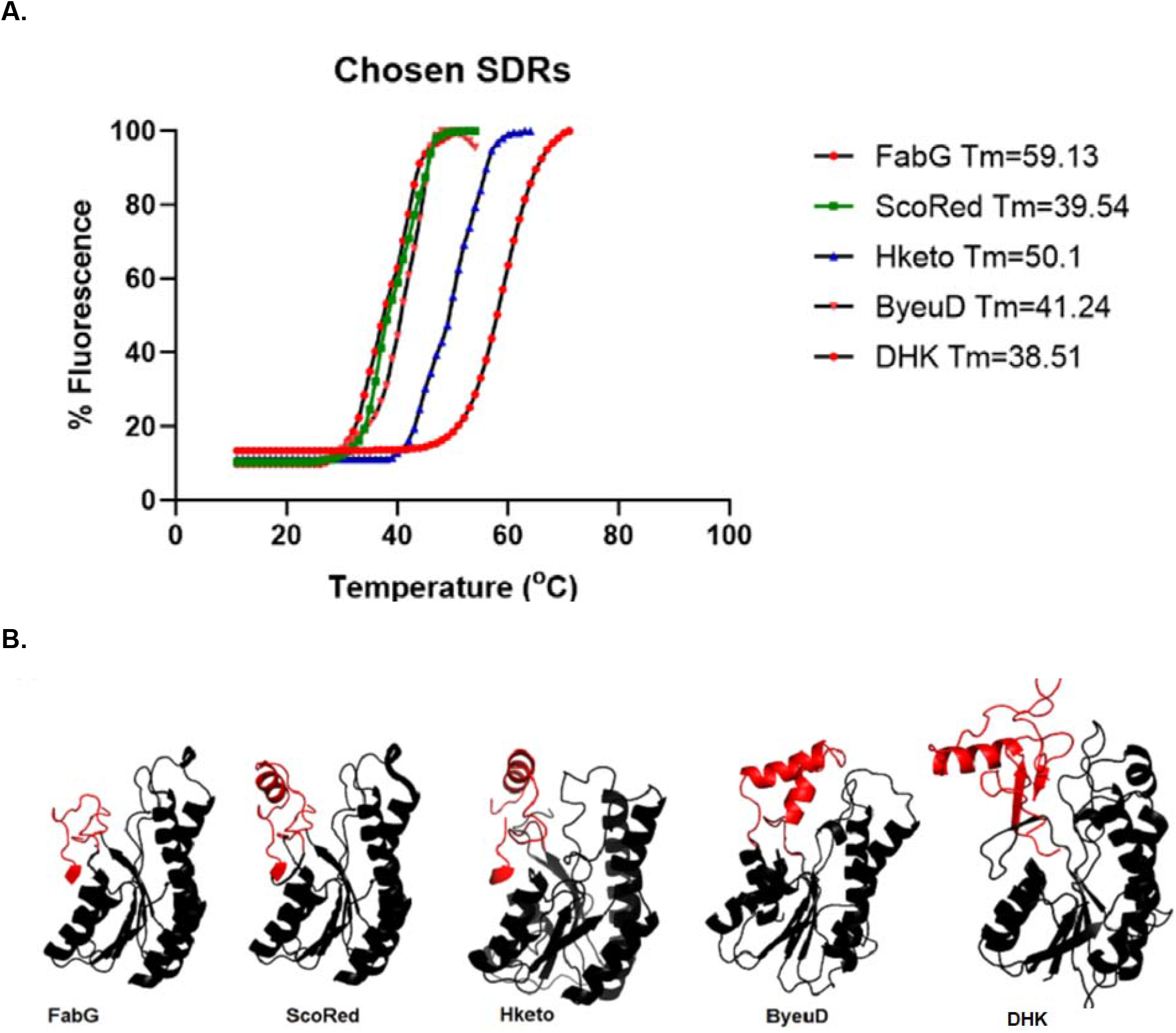
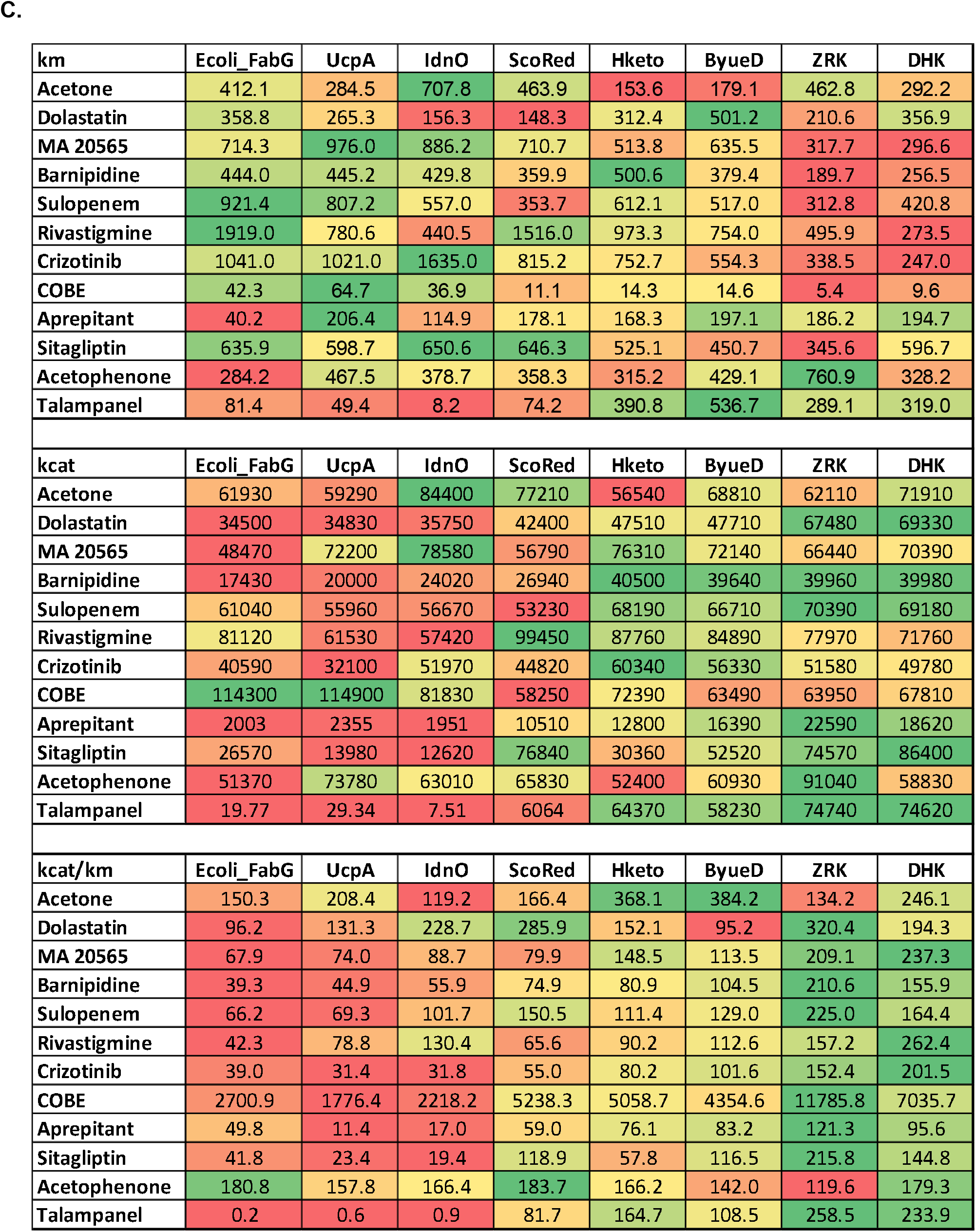
Relationship between flexibility and activity of the data mined SDRs A. Differential scanning fluorimetry of Short-chain dehydrogenases. B. The increasing complexity of the Lid-loop structure (red) of chosen SDRs. C. Heat map of the catalytic efficiency of the datamined SDRs Units: km=uM kcat=uM/min.

The DSF experiments exhibited *Synechococcus* FabG to have a higher melting temperature than the rest of the SDRs. Meanwhile, DHK has a similar melting temperature as ZRK at the lower end, followed by ByueD and Hketo, respectively. Conversely, ScoRed had a Tm like Hketo and ByueD despite having a smaller lid-loop structure as FabG (Figure 6A and 6B). Once again, the DSF data shows a direct correlation between the Tm and the lid-loop structure in SDRs (Figure 6A and 6B).

Enzyme kinetics was performed against substrates consisting of commonly published ketones such as acetophenone, α-chloroacetophenone and Ethyl-4-chloro acetoacetate. Further, other pharmaceutical ketone intermediates (K-INT) such as Talampanel K-INT, Aprepitant K-INT, Sitagliptin K-INT, Rivastigmine K-INT, Dolastatin K-INT, Sulopenem K-INT, MA-20565 K-INT and Crizotinib K-INT useful in drug industries were introduced as well (Supplementary figure S1).

Enzyme kinetics data reveal that *Synechococcus* FabG has the highest km and low kcat (thereby having the highest catalytic efficiency or kcat/km ratio except for Eco_FabG) for most of the substrates. Comparatively, ScoRed is better than previously analysed UcpA and IdnO. Hketo has a slightly larger lid-loop structure compared to those above low-molecular-weight classical SDRs. It depicts a better catalytic efficiency despite being clustered as a low-promiscuous candidate. Comparatively, ByueD has a larger lid-loop and has better catalytic efficacy than previous enzymes. However, DHK with a similar structure and a larger lid-loop portray low km with high catalytic efficiency than other SDRs. Interestingly, ScoRed exhibited the best kcat/km values for Dolastatin K-INT and Acetophenone whereas; FabG showed comparably good activity against acetophenone, α-chloroacetophenone and Ethyl-4-chloro acetoacetate (COBE). Nevertheless, overall, ZRK and DHK proved superior for the catalytic conversion of industrially relevant pharmaceutical intermediates (Figure 6C).

The standard deviations of in vitro enzyme kinetics (Figure 7A) show that they do not vary as much as the catalytic efficiency. This lack of variation indirectly depicts the homogeneity of the active site among all the SDRs (Figure 7B). The extended SDRs are more flexible as lesser variation is present among the substrates. This characteristic is especially true for ZRK and DHK.

**Figure 7:**
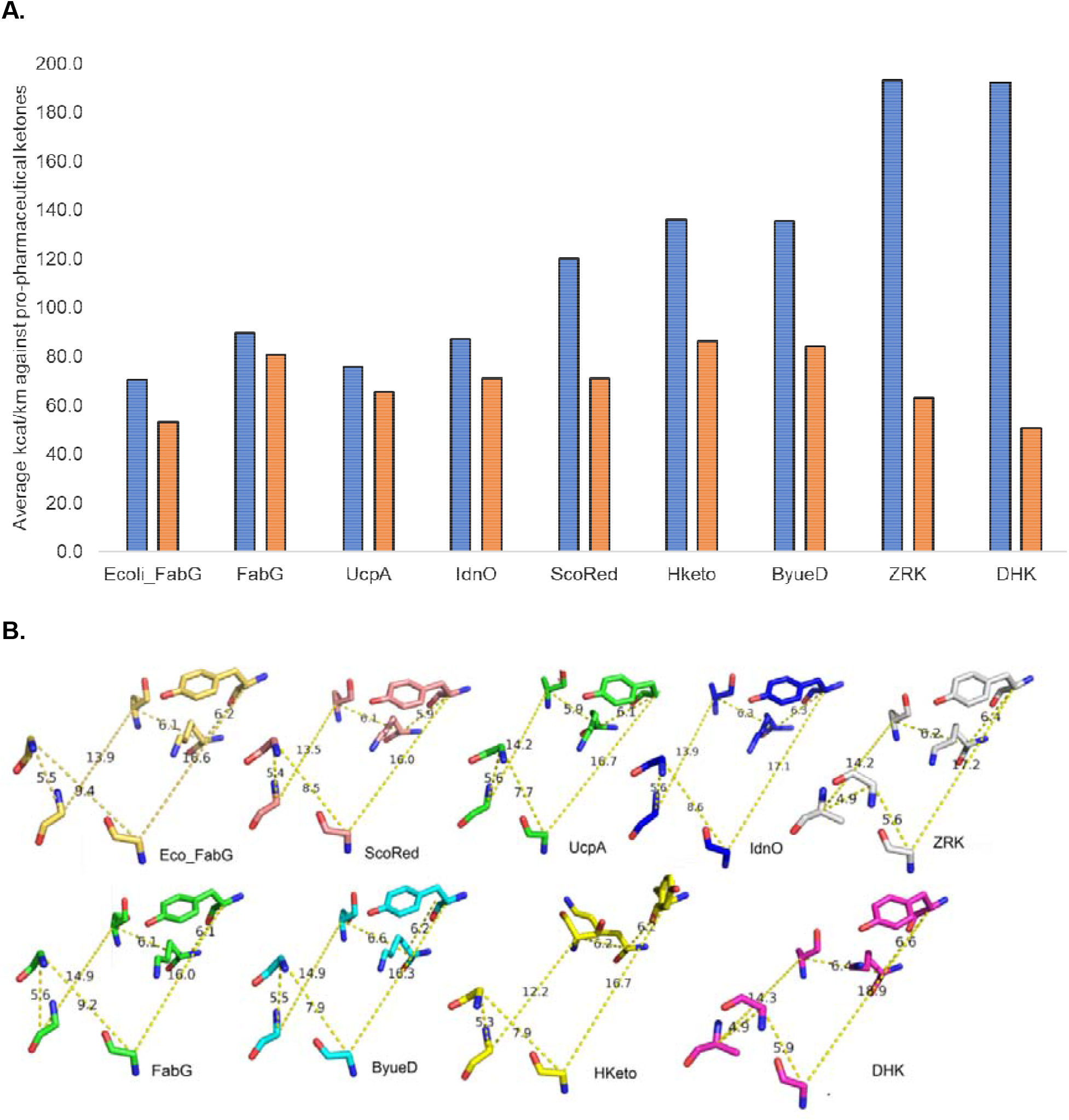
A. Average catalytic efficiency of ketoreductases with their respective standard deviations. The graph depicts an indirect measure of the flexibility of the active site as the change is quite similar to each other despite the large difference in catalytic efficiency B. The geometry of the active site and substrate-binding site between the chosen Short-chain dehydrogenases shows a similarity in the distances between the participating residues.

## Discussion

Many studies demonstrate the production of chiral alcohols, but only a few have prevailed in making any economic advances in this field. Invariably, there are hurdles in obtaining an enzyme that can perform a function of interest because the substrates are non-native. Besides, a more straightforward method can alleviate the cost and time-intensive techniques like high throughput screening and genome/metagenome sequencing. However, the onus of being a catalyst in an industrial setup with low-cost input is a tough challenge and is necessary for a large-scale setup (32)(33). Additionally, finding a promiscuous enzyme is a superior alternative as it avoids protein engineering and high throughput screening. Therefore, using machine learning for unearthing a promiscuous candidate is a favourable solution in this framework.

The modelling results show a change in protein architecture due to the variations in the lid-loop structure (Figure 2). Biochemically, classical SDRs have evolutionarily pre-existed their extended counterparts as they are critical players in essential biochemical pathways (34). Therefore, classical SDRs were modelled to architecturally link them to the ‘Extended’ class of proteins. Although connecting them in terms of evolution is an arduous task, SDRs (especially from microbes and unicellular eukaryotes) have a conserved N-terminal structure and a variable C-terminus substrate interacting region that gives a perspective on the change in the architecture of SDRs as a whole. Also, Extended SDRs are naturally promiscuous contenders for detoxification and generation of secondary metabolites in yeast and higher eukaryotes. Therefore, this superfamily is an essential placeholder for ascertaining promiscuous ketoreductases.

The lid-loop structure of the short-chain dehydrogenases is speculated to participate in the entry and exit of the substrate (35). The structure stabilizes upon the binding of NADPH and helps maintain the protein’s overall architecture (36). It has also been altered using site-directed mutagenesis for attaining a specific product of interest, e.g., the lid-loop FabG in *Synechococcus* sp. was engineered to mimic the flexibility of another native SDR, the PhaB, for increasing the production of Polyhydroxy butyrate. Conversely, the flexibility was reduced in *Serratia marcescens* SDR for the production of (R)-phenylephrine (37)(38). Contextually, it is possible to engineer SDRs to obtain a specific product of interest; however, for multi-substrate catalysis, the overall architecture, flexibility, and an estimate of catalytic activity need to be considered for establishing a viable candidate. Therefore, firstly, it is crucial to establish the relationship between flexibility and catalytic efficiency. Secondly, enzymes belonging to a similar group with comparable physicochemical properties with diverse lid-loop sizes are favourable for locating a sustainable enzyme.

To prove the first assumption, the metabolically essential SDR Eco_FabG (*E. coli* FabG) and other non-essential SDRs (IdnO and UcpA) with similar identity demonstrate a clear relationship between flexibility and catalytic efficiency (Figure 3A). Also, the general direction for prospecting a multi-substrate catalyzing SDR could be their non-essentiality. Therefore, a reasonably well established, non-essential SDR from ZRK was chosen as a positive control for this experiment. It demonstrated the relationship between flexibility and catalytic efficiency for multiple substrates (Figure 3C). Also, modelling data showed ZRK, an extended class SDR, to have a larger lid-loop (Figure 3B).

This size extension could contribute to the overall flexibility of the protein as well. Therefore, for the second assumption, we focused on sorting the proteins based on their physicochemical properties. To this end, we used machine learning to classify and cluster enzymes similar to reported promiscuous candidates. We also chose candidates with different molecular weights from FabG (∼250 residues) to ZRK (∼320 residues) for forming a spectrum of non-essential enzymes with increasing lid-loop architecture and expectantly flexibility.

Deep learning and machine learning have been used to gain a better perspective on protein structure and function. Also, protein sequences are a good metric for predicting the kingdom of origin and thermostability (39)(40). Both systems have advanced quite a lot in the last decade, e.g., Chakravorty *et al*. used structural data, protein, the encoding nucleotide sequence as features for classifying thermostable and mesothermal proteins. Many unsupervised and supervised methods were used to this end to determine the cause of thermostability at ‘each level’ (i.e., nucleotide, protein, structure level). The study showed that protein sequence data is the best measure for understanding protein thermostability with 90.97% accuracy (41).

Similarly, classification of protein stability using network features (in addition to structural and amino acid compositions) using network centrality values and sequence features (specifically di- and tri-peptide bonds) was quite successful. The centrality depends on the distances between Cα atoms that correlate better with protein stability. These features improved classification significantly to 96% accuracy (42).

A processed dataset exemplifies the accuracy and performance of the algorithms in question. Therefore, we refined the dataset by considering the gradual change in genomes (and subsequently proteins) influenced by speciation. Genes (and proteins, by extension) evolve from gene duplication and mutation events (43). Compared to these two occurrences, gene insertions are rare. Genomic studies across the three domains of life have shown that successful gene shuffling has occurred only 48 times in higher eukaryotes (i.e., between *Caenorhabditis and Drosophila*) in half a billion years. Also, gene shuffling is one order of magnitude lesser than gene duplication in eukaryotes, unlike prokaryotes, where both incidences are similar. Therefore, this shows that the introns are not necessarily involved in gene shuffling as prokaryotes lack them and yet, gene shuffling is a common artefact among them (44). Therefore, SDRs were classified based on their phylum of origin. Also, as mentioned earlier, features based on protein sequences are superlative for predicting thermostability, an invariable measure of flexibility. Classification effectively removed the outliers (of evolution) and gave a refined dataset of true positives, which was further clustered to provide unknown candidates closer to the reported promiscuous members.

The clustered true positives (Hketo, ByueD, DHK) lying close to the reported mesophilic promiscuous proteins were selected. Also, ScoRed and *Synechococcus* FabG were chosen for establishing the in vivo capability of reported candidates similar to ZRK. Moreover, the desired enzymes were modelled to discern their lid-loop structures (Figure 6B). The DSF assay exhibited that lid-loop size is inversely proportional to the melting temperature. ScoRed is an exception as it showed a similar melting temperature as Hketo, ByueD and DHK (Figure 6A). However, the catalytic efficiency of all the enzymes established that lid-loop is an essential component for instituting flexibility and promiscuity in SDRs. The large lid-loop containing DHK and ZRK are superior for converting pharmaceutical ketone intermediates. Further, Hketo and ByueD have the next best choice catalytic efficiency (Figure 6C). Although *Synechococcus* FabG and ScoRed have good kcat/km values, they are relatively less than the SDRs with larger lid-loop.

The clustered true positives (Hketo, ByueD, DHK) lying close to the reported mesophilic promiscuous proteins were selected. Also, ScoRed and *Synechococcus* FabG were chosen for establishing the in vivo capability of reported candidates similar to ZRK. Moreover, the desired enzymes were modelled to discern their lid-loop structures as well (Figure 6B). The DSF assay exhibited that lid-loop size is inversely proportional to the melting temperature. ScoRed is an exception as it showed a similar melting temperature as Hketo, ByueD and DHK (Figure 6A). However, the catalytic efficiency of all the enzymes established that lid-loop is an essential component for instituting flexibility and promiscuity in SDRs. The large lid-loop containing DHK and ZRK are superior for converting pharmaceutical ketone intermediates. Further, Hketo and ByueD have the next best catalytic efficiency (Figure 6C). Although *Synechococcus* FabG and ScoRed have good kcat/km values, they are relatively less than the SDRs with larger lid-loop.

It is critical to understand that engineering proteins are subtle, and an angstrom difference in the right places in an enzyme can be a ‘make or break’ for catalysis (1)(45). Therefore, a strategy to identify a potential SDR based on flexibility and architecture is a less expensive and valuable strategy for converting multiple ketones of pharmaceutical importance. Further, it leads to costly high throughput screening as well. More than ∼100,000 (and counting) SDRs and machine learning of primary sequences can help detect attractive test candidates.

## Author contributions

SD and RS conceived and provided support via a grant for the study. APS, SR and AG performed all the preliminary and final experiments. NK was involved in designing and executing the in vitro and modelling experiments. All authors have read and approved the final manuscript.

## Acknowledgement

The financial support from the Biotechnology Industry Partnership Program, Department of Biotechnology, India (DBT Sanction Order No: BT/BIP0475/13/11) and Grant BT/PR13429/BRB/10/767/2009 to Prof. S. Ramaswamy is gratefully acknowledged.

## Methodology

### 1. Reagents and chemicals

A set of high value prochiral ketone intermediates namely, Talampanel ketone intermediate or K-INT (3,4-methylenedioxyphenyl acetone), Barnipidine K-INT (1-N-carbobenzoxy-3-pyrrolidione), Aprepitant K-INT (1- (3,5-bis-trifluoromethyl-phenyl)-ethanone), Crizotinib K-INT (1- (2,6-dichloro-3-fluoro-phenyl)-ethanone, 3-trifluoromethyl acetophenone), Dolastatin K-INT (2-phenyl-1-thiazol-2-yl-ethanone, 1- (3-methoxyphenyl) ethanone), Sitagliptin K-INT (3-oxo-4- (2,4,5-trifluoro-phenyl)-butyric acid methyl ester) and Rivastigmine K-INT (1- (3-methoxyphenyl) ethanone) were synthesized as reported elsewhere (46). The uses of these pharmaceutical intermediates are highlighted in Supplementary Figure 1.

### 2. Homology modeling

The homologs for the enzymes of interest were identified using Protein BLAST. The best hit was used as a template for homology modelling using SWISS-MODEL webserver and was validated by PROCHECK software. Further, the obtained model was then refined using AlphaFold through ChimeraX interface.

### 3. Flexibility prediction

Further, the flexibility of the modelled proteins was checked by using B-factor in ChimeraX. Furthermore, tools for predicting protein flexibility namely, Medusa (a server which uses deep learning to predict protein flexibility), PredyFlexy and FlexServ were used for validating the AlphaFold models (47)(48)(48).

### 4. Cloning of the genes

- The genes native to *Escherichia coli* viz. Ecoli_FabG, IdnO and UcpA were acquired from the ASKA collection. The enzymes are cloned in pCA24N plasmid which has T5 promoter which is compatible with T7 polymerase in BL21(DE3) strain (49).
- Genes encoding the other heterogenous enzymes namely, *Sulfolobus solfataricus* alcohol dehydrogenase (SSADH), *Zygosaccharomyces rouxii* SDR (ZRK), *Hansenula polymorpha* DL-1 peroxisomal 2,4-dienoyl-CoA reductase (Hketo), *Synechococcus sp*. PCC 7942 3-ketoacyl-[acyl-carrier-protein] reductase (FabG), *Bacillus sp*. ECU0013 ADH (ByueD) and *Debaryomyces hansenii* Ketoreductase (DHK) were codon optimized for *E. coli* and synthesized by Geneart (http://www.geneart.com).
- The gene sequences are deposited in DDBJ and Genbank with the following accession numbers namely, LC325171 for SSADH, LC325172 for ZRK, LC325173 for PAR, LC325174 for ByueD, LC325175 for Hketo, LC325176 for FabG, XP_458533.2 for DHK.
- These are then sub-cloned into pET28a vector (Novagen) using the restriction sites NdeI and XhoI with an N-terminal His-tag in frame.

### 5. Purification of protein

- The cells are harvested by centrifugation (8000g ×20 min) and resuspended in lysis buffer (50 mM Sodium Phosphate Dibasic, pH 8.0, 0.3 M NaCl, 10 mM imidazole). The cells are lysed on ice by sonication at 39% amplitude with ON and OFF cycle of 10 seconds for 7 rounds and the debris removed by centrifugation (22000 g × 30 min).
- The recombinant His tagged protein is purified by Ni-NTA affinity chromatography (4ml, Qiagen) and eluted by using elution buffer (50 mM Sodium Phosphate Dibasic, pH 8.0, 0.3 M NaCl, 250 mM imidazole).
- The protein is concentrated to 10 mg/mL with 10,000 MWCO centricons (Millipore). To remove imidazole the protein is diluted in 20mM Sodium Phosphate buffer pH 7.2 and concentrated again. This procedure is repeated 3 times until ∼0.25mM imidazole remained.
- Soluble protein expression experiments are performed as in the protocol followed for whole-cell transformation. The cells are harvested 18h post induction and the soluble His-tagged ketoreductases are purified by Ni–NTA affinity chromatography.
- The SDS PAGE of the purified enzymes is depicted in Supplementary figure S2.

### 6. Buffer optimization and Thermal shift assay (TSA)

- The purified protein is obtained from -80°C stock (prepared after purification) and thawed in ice for ∼30 minutes. The protein concentration is measured in a spectrophotometer (A_215_/A_225_) and the protein is diluted to 1mg/ml.
- For a single reaction, 50ug/ml of final protein concentration is used along with 20x SYPRO orange (Final concentration). The reaction is run in an RT PCR machine (Bio-Rad/Applied Biosystems 7900HT) at 2% ramp rate (∼1°C per minute) and from 25°C to 90°C at Ex/Em (497/520).
- The run is approximately 40 minutes. The raw data is extracted into Graphpad Prism and the Tm (V_50_) is calculated using Boltzmann sigmoidal fit to obtain the melting temperature of protein.

### 7. In Vitro Kinetic Assays

- Enzyme specific activity

The measurement of exam activity is done in the Tecan S200 spectrophotometer at A_340_. The final concentration of enzyme used is 1ug/ml for the reaction. The substrate is added to approximately 200 times the concentration of the enzyme. A consecutive rate of equal conversion is used to establish the specific activity of the enzyme. It is then extrapolated for 1uM/min/mg (specific activity).

- Enzyme Kinetics

For ketoreduction, all assays are performed in optimal buffer conditions with the enzyme concentration at 1ug/ml. The km for each ketone of interest is obtained by varying the substrate concentration in the presence of 2000μM NADPH/NADH. The Km for NADPH/NADH is obtained by varying the cofactor concentration in the presence of saturating amounts of ketone substrate. The slope for initial steady-state reaction is used for obtaining the velocity (uM/s) at known substrate concentrations. These two parameters are plotted in GraphPad Prism to obtain a Michaelis Menten fit to obtain Km, Vmax and Kcat of an enzyme for a substrate of interest.

### 8. Data extraction

#### (i) Data collection

A domain specific search is done on Uniprot server to extract all the data for SDR. Also sdr-enzymes.org, an HMM based classification website, was used to gather all the protein sequences. The data added in MS-Excel spreadsheet and redundancy of the sequences of removed using CD-HIT server with sequence identity cut off as 0.9 and also, conditional formatting in Excel was used to verify the same. Physicochemical parameters: NCBI taxonomy identifier, Algabase, Bacterio.net, Encyclopedia of Life (EOL) databases were used to identify the Species, Phylum and Environment of survival. These were added individually in the database.

ProtParam and Protscale were used to measure Total Amino acids, Enzyme name, Sequence, Various Amino Acids %, Molecular Weight, Theoretical PI, GRAVY (Hydrophobicity), Extinction coefficient, Aliphatic Index, Negative %, Positive % and Solubility were used as features for classifying proteins.

Normalization: Transformation of the data to a homogeneous and final dataset was performed applying the following equation:

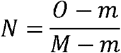

Where (N) denotes normalized output, O represents the original data value, M and m refers to the maximum and minimum data values of a parameter, respectively.

### (ii) Machine Learning: Feature selection, Classification and Clustering

Scikit, a free machine learning and clustering (Scipy) tool based on Python programming language, which allows machine learning and clustering through cluster-computing and iterative processes, was used for finding.

### (iii) Supervised learning methods

In supervised learning methods, part of the dataset is used for training the system to recognize hidden patterns and classify them accordingly. In this exercise, the system was trained using 90% of the dataset and 10% was used for testing. This was iterated multiple times in order to get a semblance of true positives. These were then used for clustering using unsupervised learning. Following are the methods used in classification.

### (iv) K-Nearest neighbour

This method utilizes distances between points as a measure for classification. It classifies the data points into different classes based on the characteristics of the nearest neighbours. The distances are measured using Minkowski, Manhattan, Cosine, Euclidean and Jaccard distance. Of these, Makowski and Cosine distances are used to measure similarities in vectors. In contrast, Jaccard’s distance is useful for smaller sample size with binary inputs e.g., DNA gel analysis. The Manhattan distance is useful for a dataset consisting of real-world geometric problems. Therefore, in the current study, Euclidean distance has been used for classifying the enzymes as it is contextually superior to Manhattan distance. The number of weights used in the current study was 3 using Euclidean distance. About 1500 iterations were carried out using Random Forest to ascertain classification.

### (v) Support Vector Machine (SVM)

The method utilizes a hyper-plane (a plane in *n* dimensions) that separates the data belonging to different classes. This is useful for classifying high dimensional data. The gamma value used for the current classification was 2.25, followed by 500 iterations.

### (vi) Random Forest (RF)

A multidimensional dataset such as the current one cannot be classified using simple binary or decision trees. Although the computation time is significantly low, the classification is poor due to the multi-dimensionality of the dataset. Therefore, the random forest (an advance on the decision trees) is better suited for the problem. It makes predictions based on consensus rather than just a binomial decision of the decision trees. The algorithm works on the principle of navigating over random subsets of features for obtaining the best feature. Hence, the search space is diversified, resulting in robust prediction. Furthermore, it can handle imbalances in sample size among the classes. About 1500 iterations were carried out using Random Forest to ascertain classification.

### (vii) Extra tree (ET)

Another algorithm that is based on decision trees is Extra trees, short for extremely randomized trees. It is an ensemble learning algorithm where, unlike random forest that develops each decision tree from a sample of the training dataset, the extra tree algorithm utilizes the complete training dataset. It works by making a large number of unpruned decision trees from the training dataset and then classifying them based on the majority. About 1500 iterations were carried out using Random Forest to ascertain classification.

### (viii) Unsupervised learning methods

In unsupervised learning the data is clustered without a primary training set. It is useful for finding low-dimensional features from a high-dimensional input data. The unsupervised learning techniques used in this thesis are as follows:

### (ix) k-means clustering

The k-means clustering is an unsupervised algorithm which is done by using a set of random numbers as *mean* denoted as ‘*k*’. After which the numbers closer in value to the *mean* are taken into consideration for forming an initial cluster. Further, the mean is continually changed until all the numbers are close to its value. The number of means depend on the number of clusters chosen for the study.

### (x) Principal component analysis

The principle component analysis (PCA) is used for reducing dimensions of the data to two. Based on the ‘*n*’ input data vectors, the PCA generates ‘*p*’ singular vectors of the data matrix.

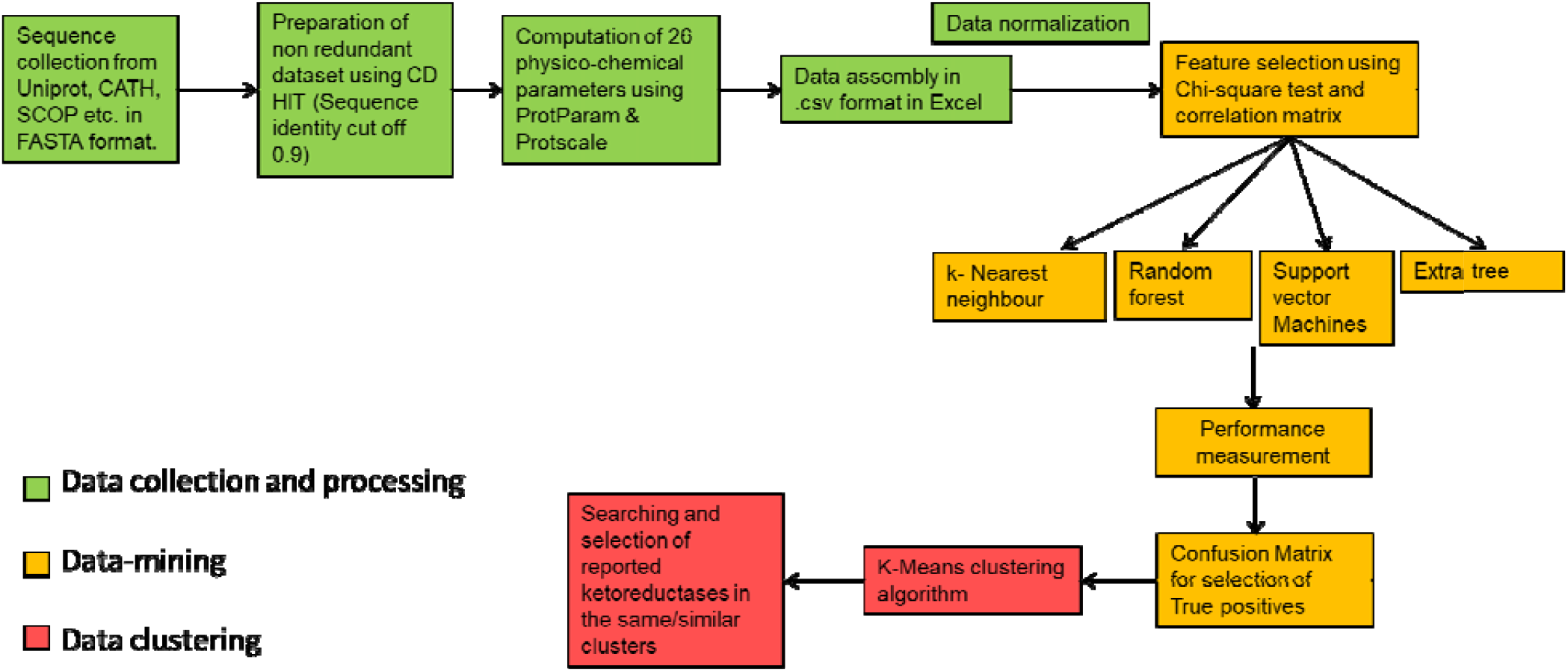
Overall schematic of obtaining proteins

