## Supplementary material for "Flexibility of Short-chain dehydrogenase is interconnected to its promiscuity for the reduction of multiple ketone intermediates": JBC_2_Supplementary.docx

**Supplementary data**

**Supplementary figure S1:** Uses of various pharmaceutical compounds derived from the chiral pro-pharmaceutical intermediates used as substrates from the repurposing of the isolated ketoreductases.

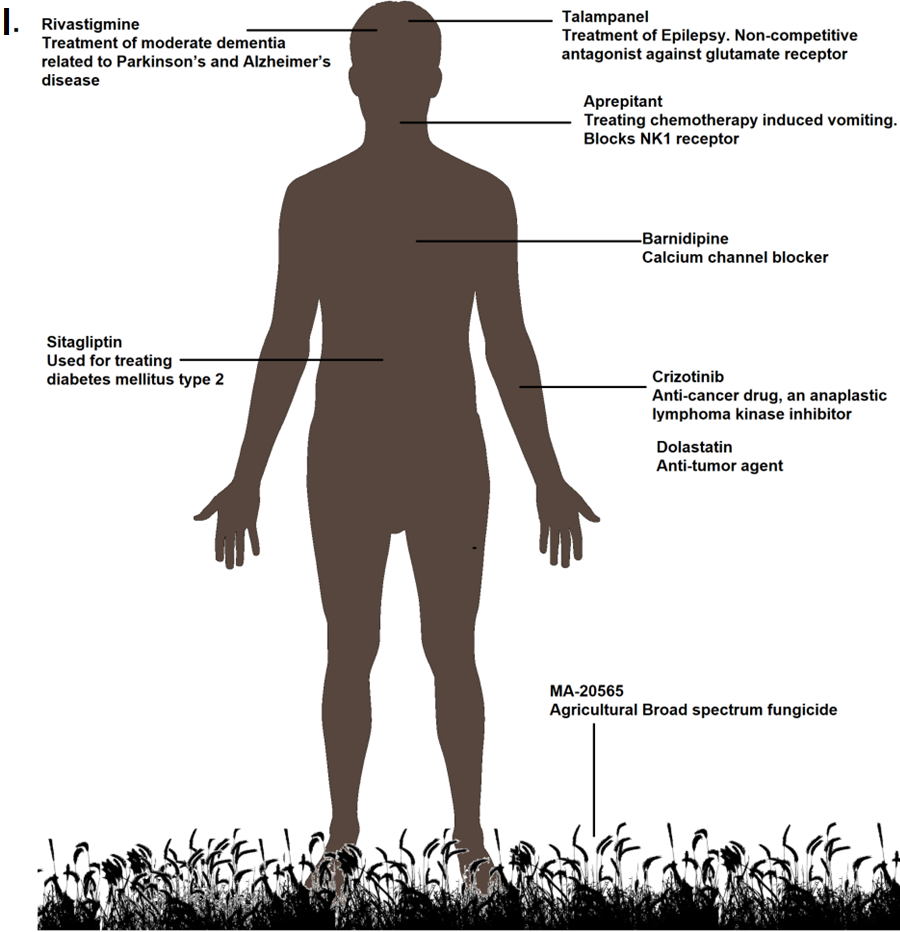

**Supplementary figure S2:** The geometry of the substrate binding site and active site GxxGxGxnSXnYxxxK) of different sized SDRs show similar dimensions (Angstroms).

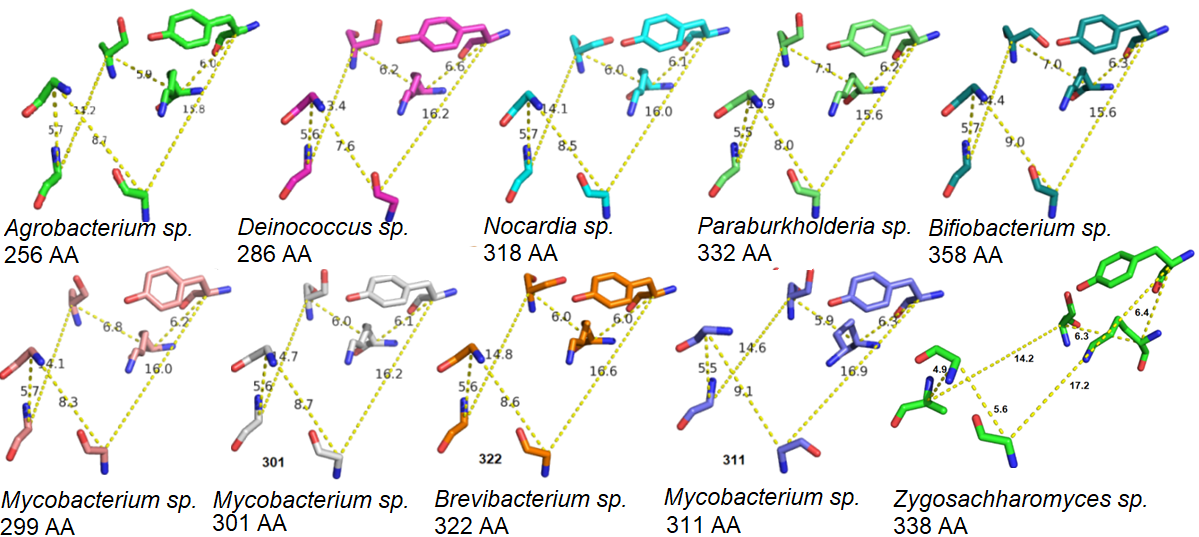

**Supplementary figure S3:** Identification and cloning of reported ketoreductases and introduction of pro-pharmaceutical ketone intermediates. Gel pictures and SDS gel pictures of isolation A. *Escherichia coli* FabG B. *Escherichia coli* UcpA (Lane 2, 3) and IdnO (Lane 4, 5) C. *Synechococcus sp.* PCC 4092 FabG D. *Debaryomyces hansenii* DHK (Lane 2) E. Lanes 1. *Zygosaccharomyces rouxii* SDR (ZRK), 2. *Hansenula polymorpha* ketoreductase (Hketo), 3. *Bacillus subtilis* yueD (ByueD), 4. 3-oxoacyl-[acyl-carrier-protein] reductase (FabG) and 5. Low range molecular weight marker.

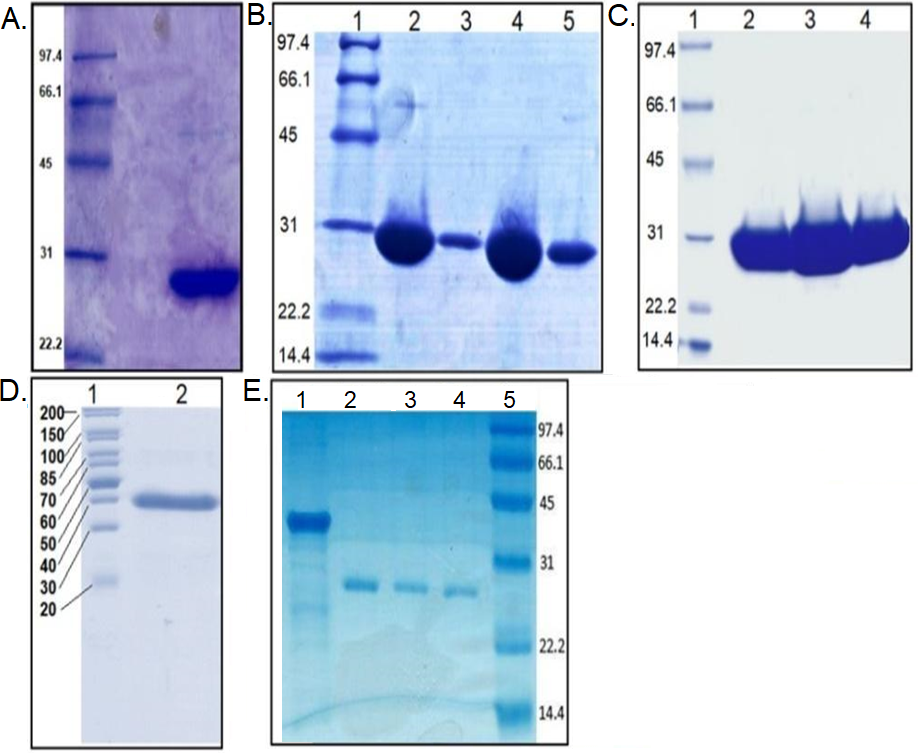

**Supplementary figure S4:**

**A.** Alpha fold models of the reported and predicted SDRs with B factor coloured from blue to red signifying low to high b factor

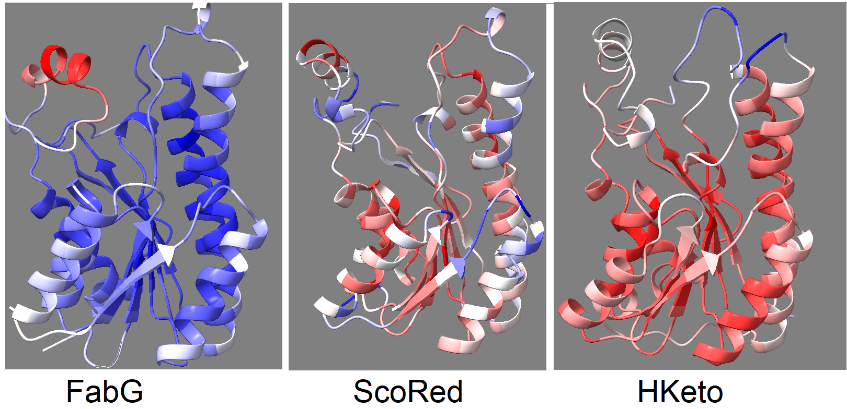

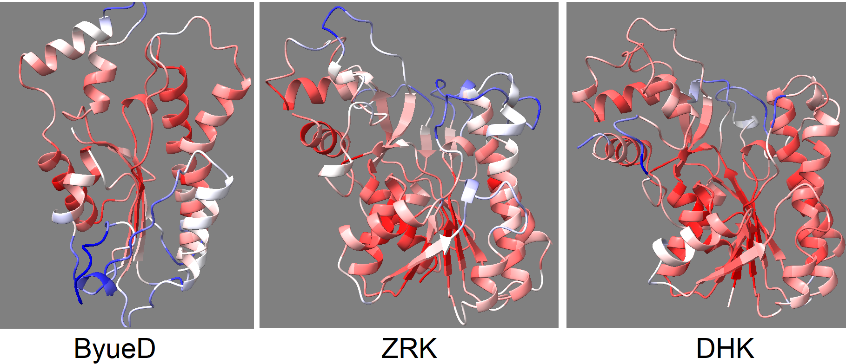

**B.** Protein flexibility predicted by Medusa depicts an increasing percentage of flexible regions in proteins having a larger lid-loop structure.

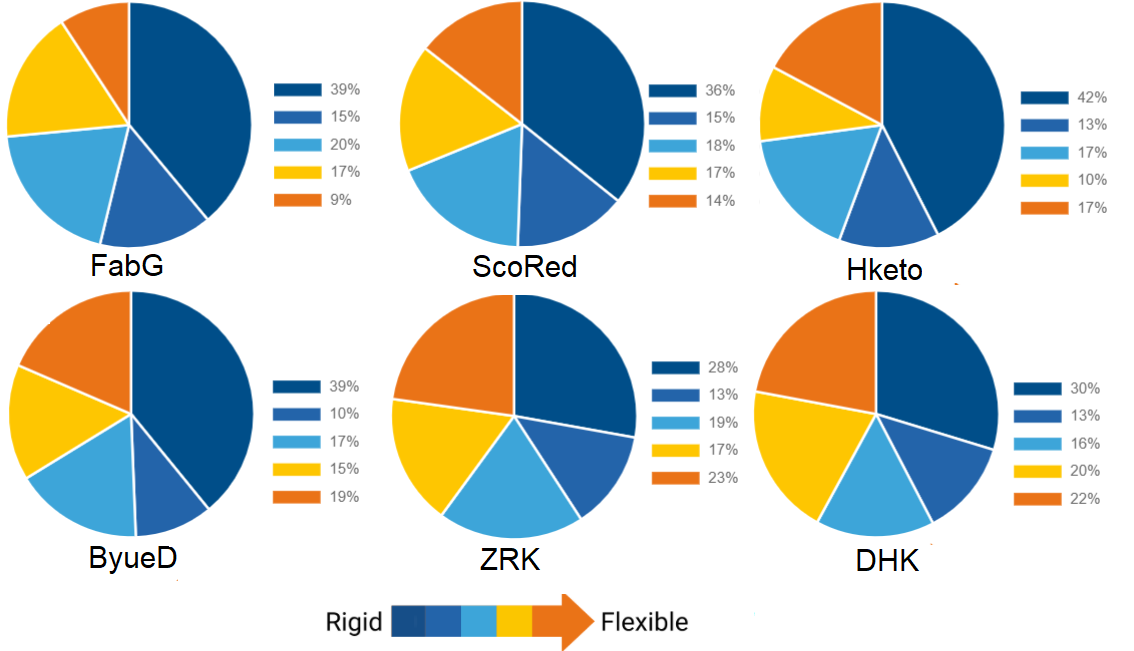

**C.** Protein flexibility using primary protein sequence using PrediFlexy server

**FabG**
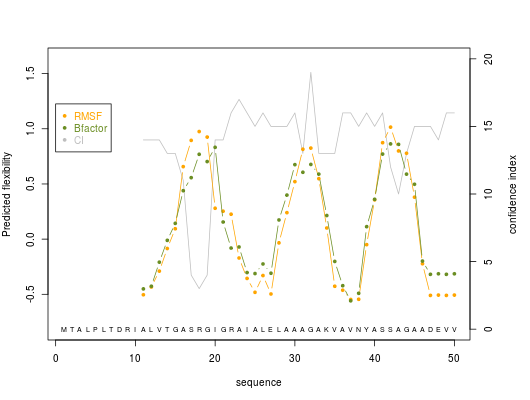
 
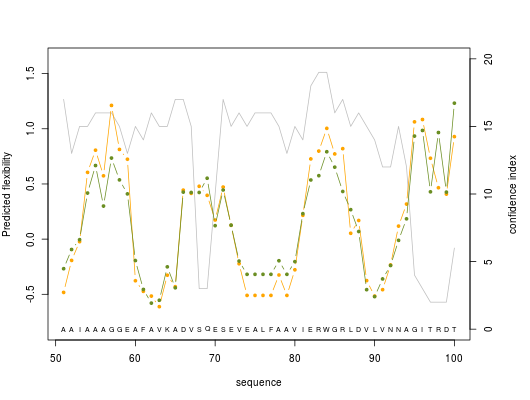
 
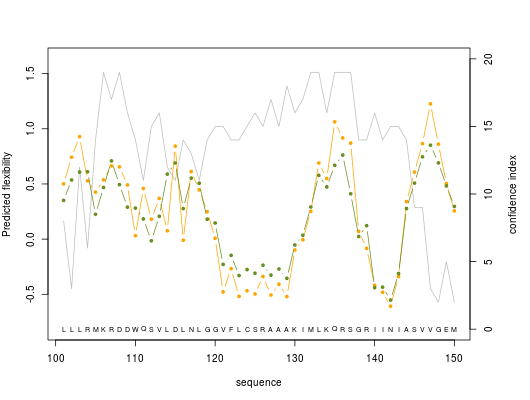
 
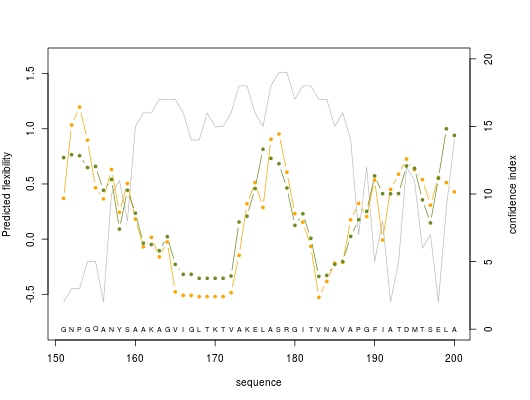
 
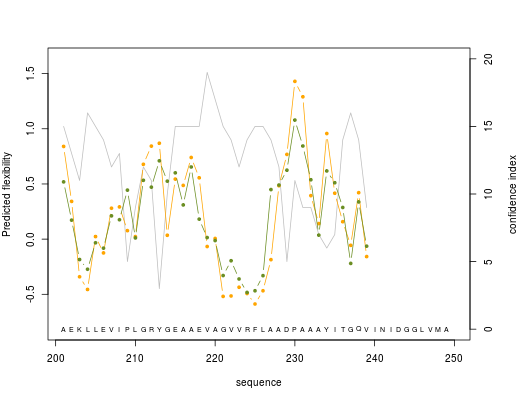

**ScoRed**

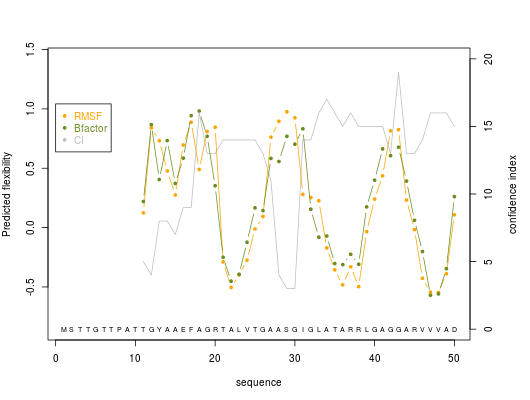
 
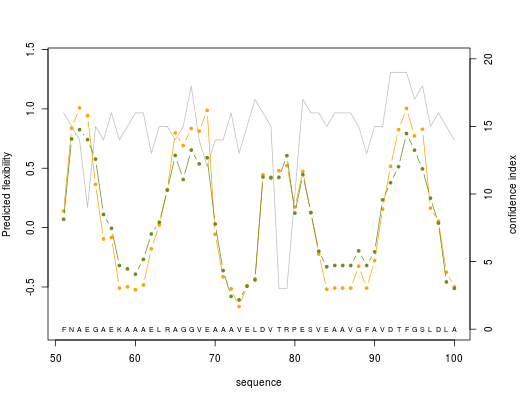
 
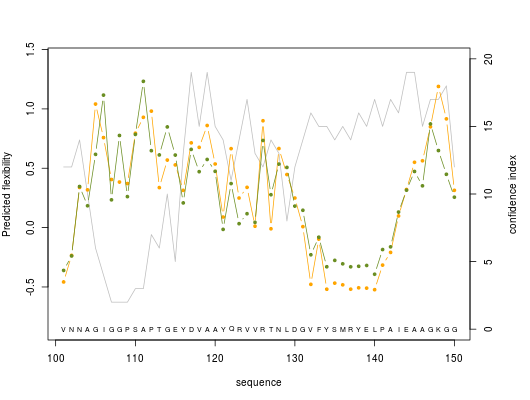
 
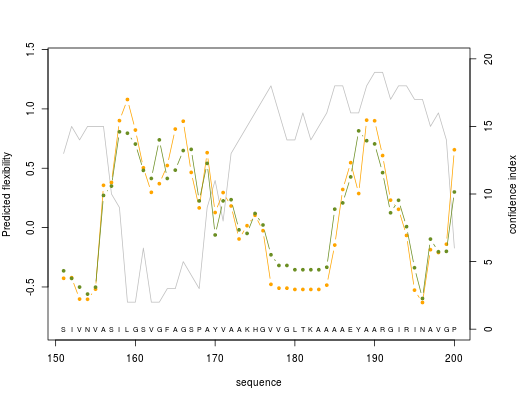
 
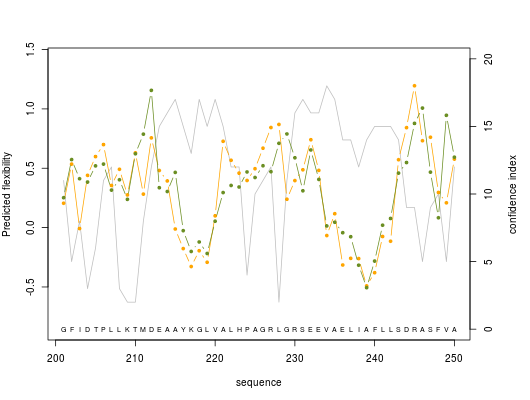
 
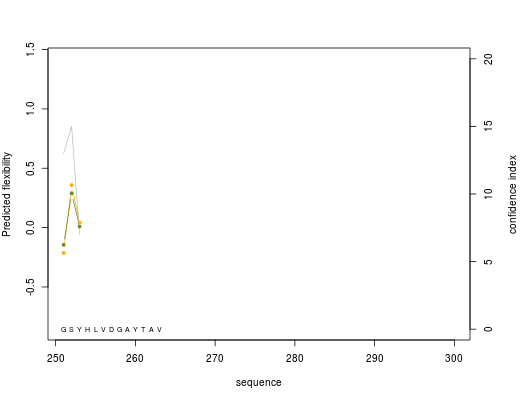

**.**

**Hketo**

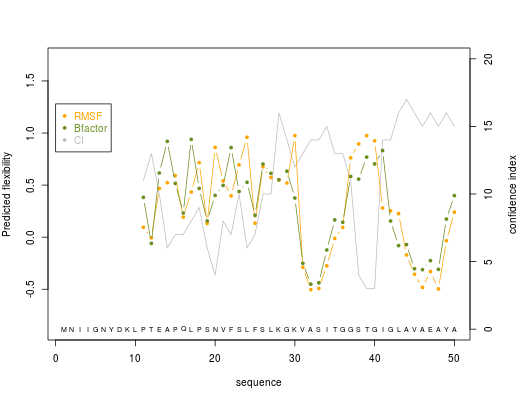
 
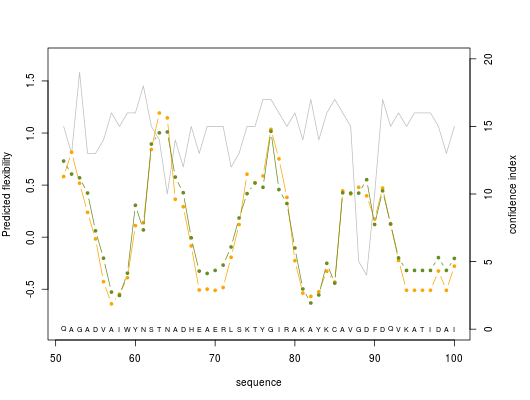
 
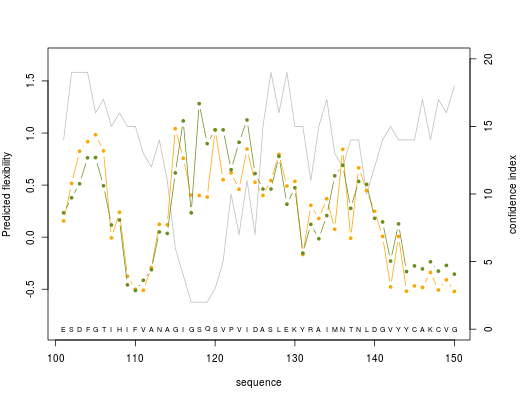
 
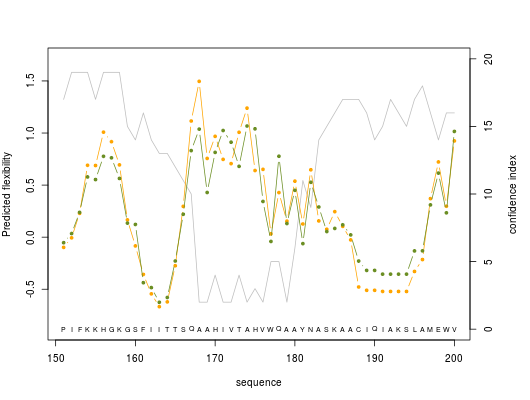
 
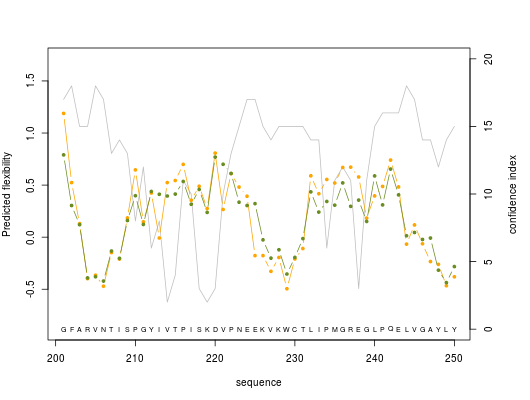
 
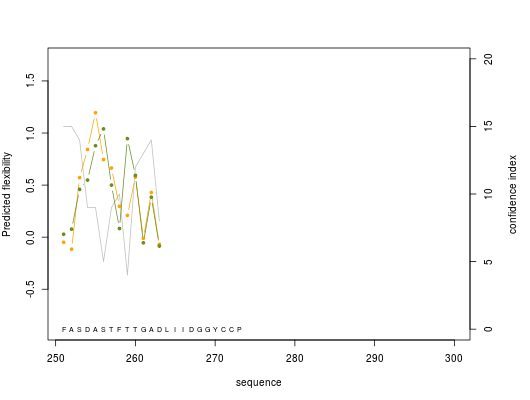

**ByueD**
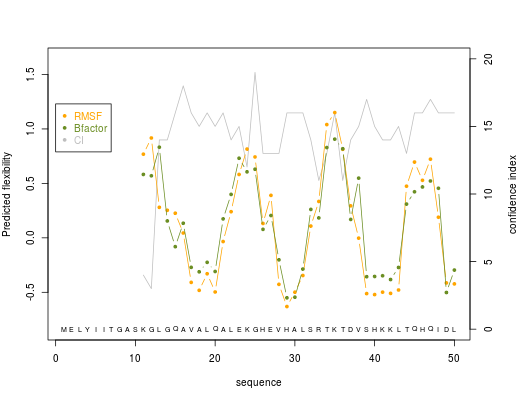
 
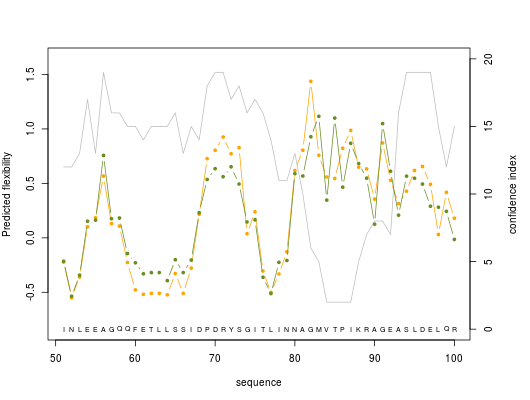
 
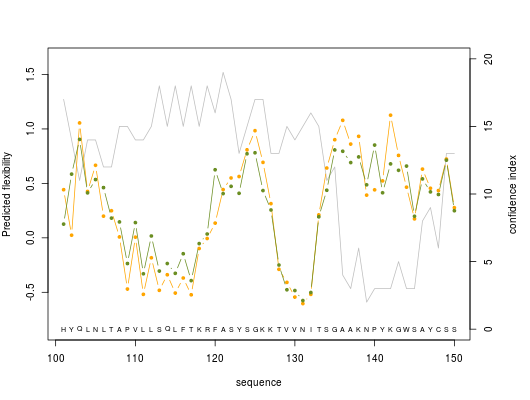
 
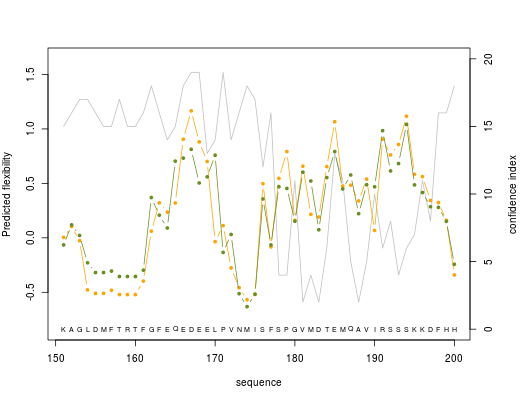
 
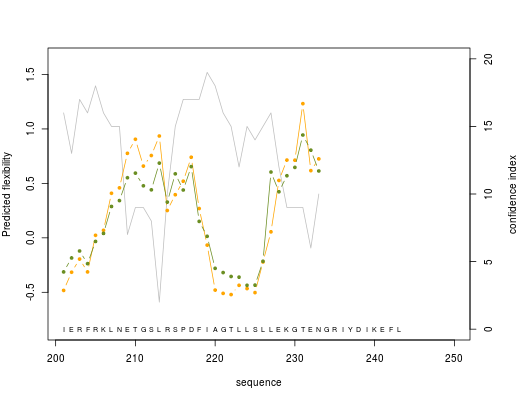

**ZRK**
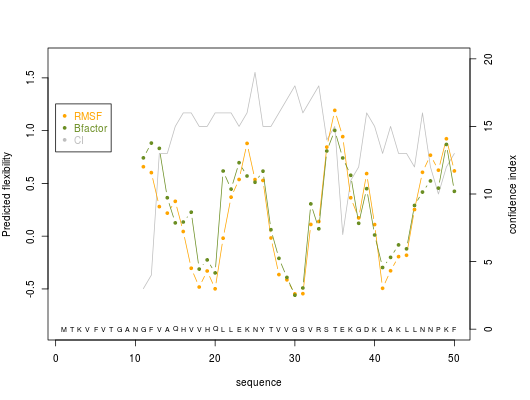
 
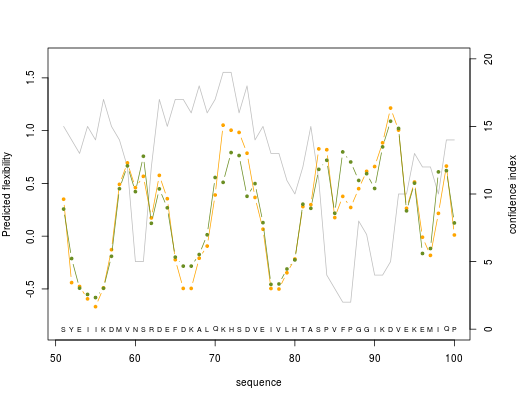
 
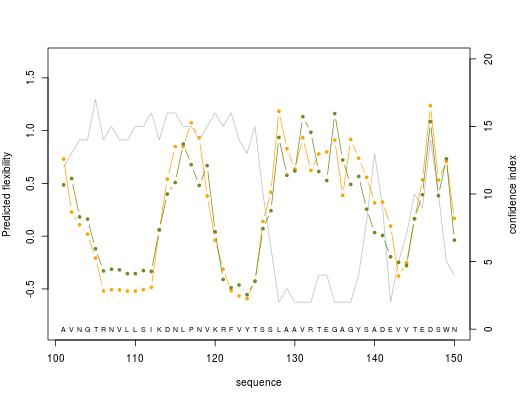
 
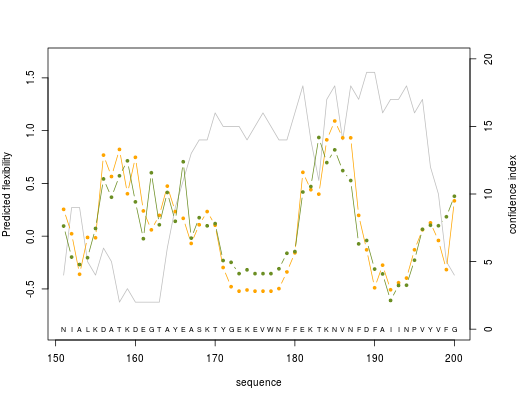
 
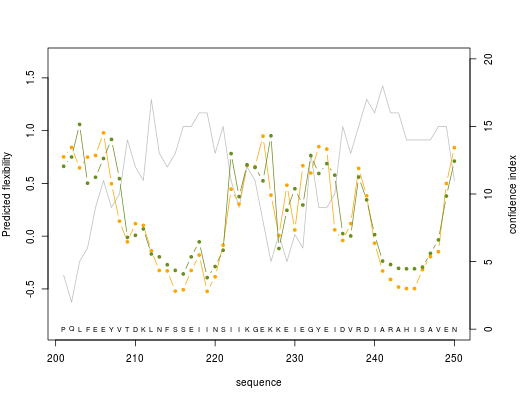
 
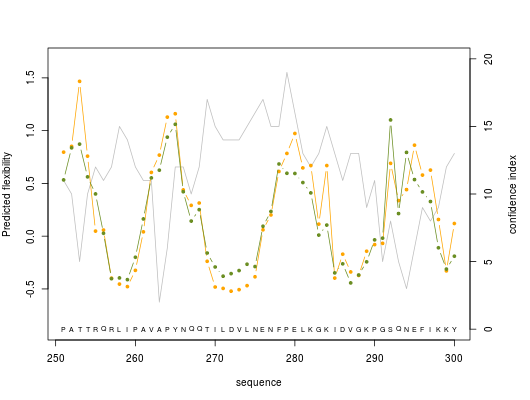
 
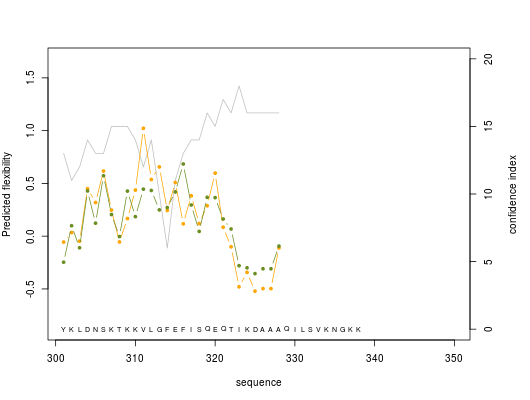

**DHK**
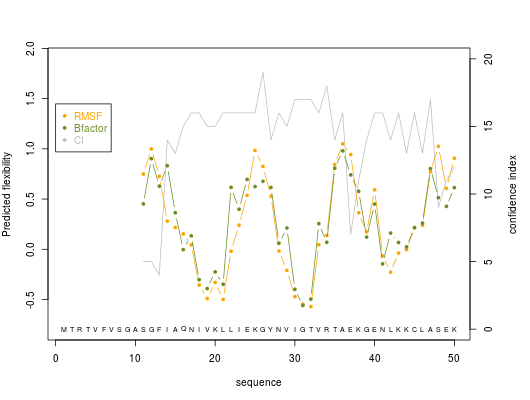
 
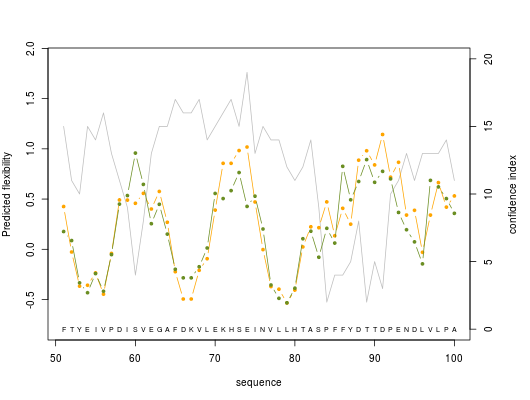
 
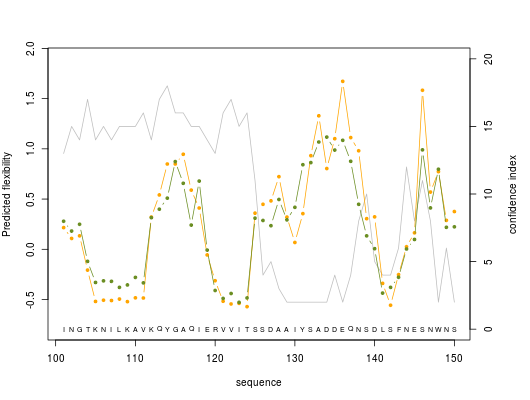
 
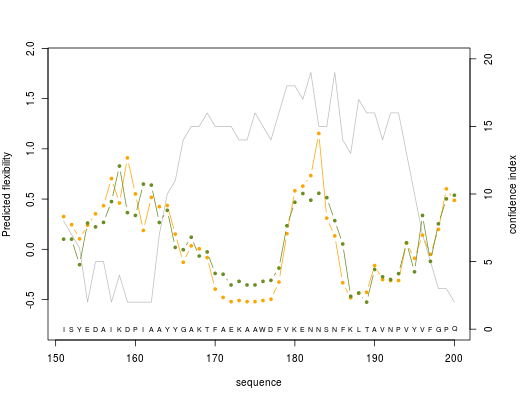
 
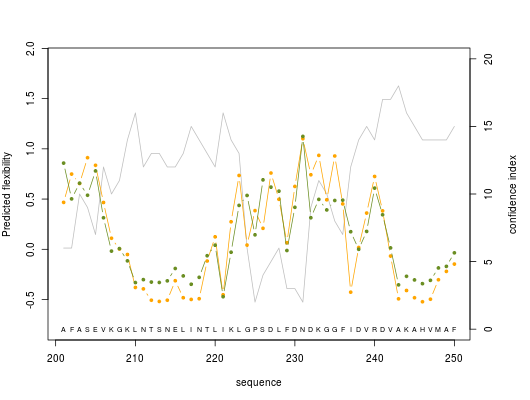
 
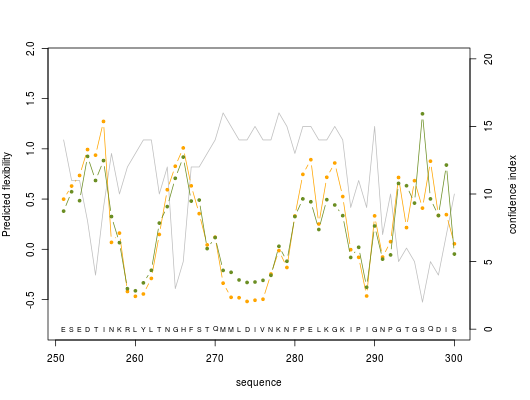
 
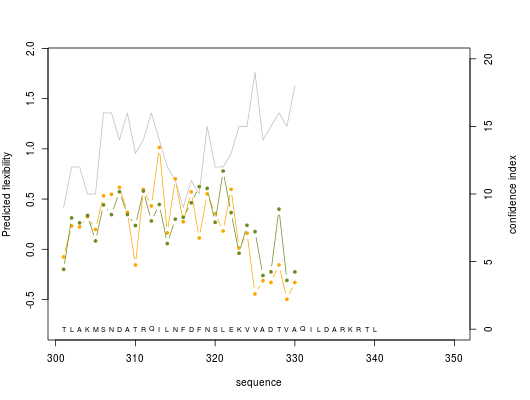

**D.** Prediction of protein flexibility using AlphaFold models using FlexServ.

**FabG**

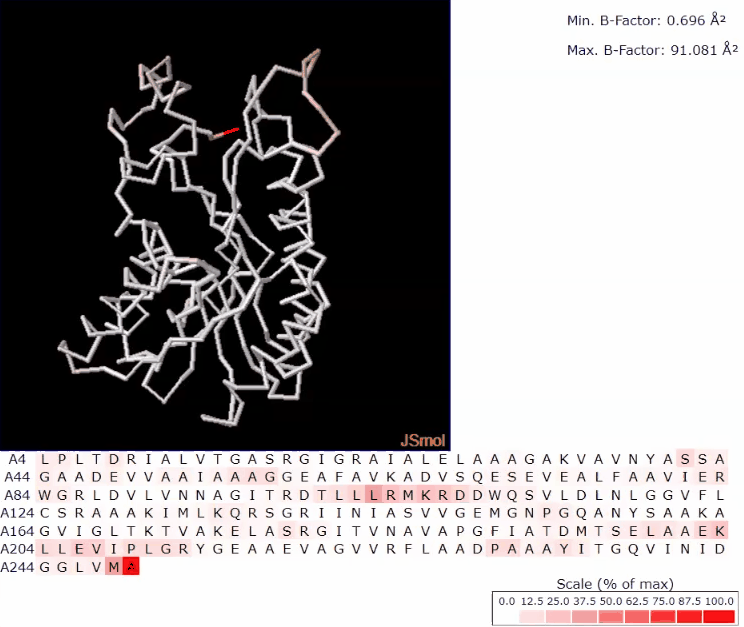

**ScoRed**

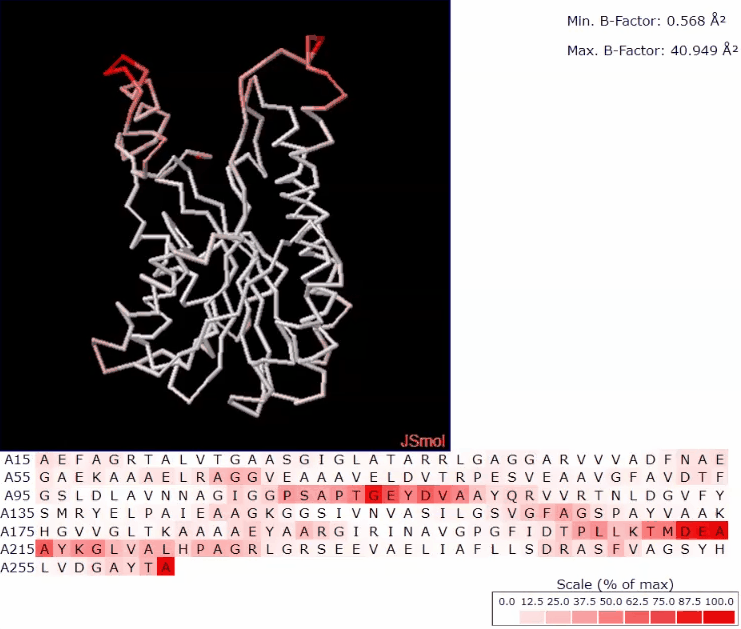

**Hketo**

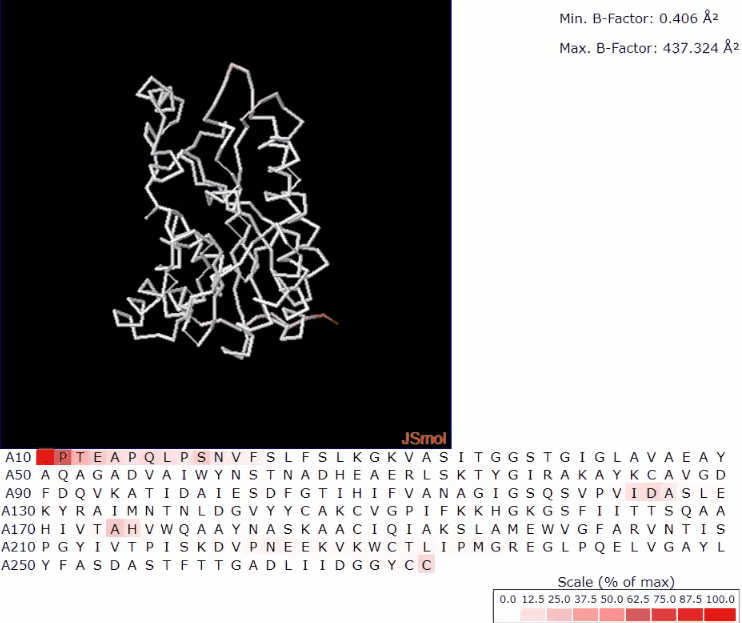

**ByueD**

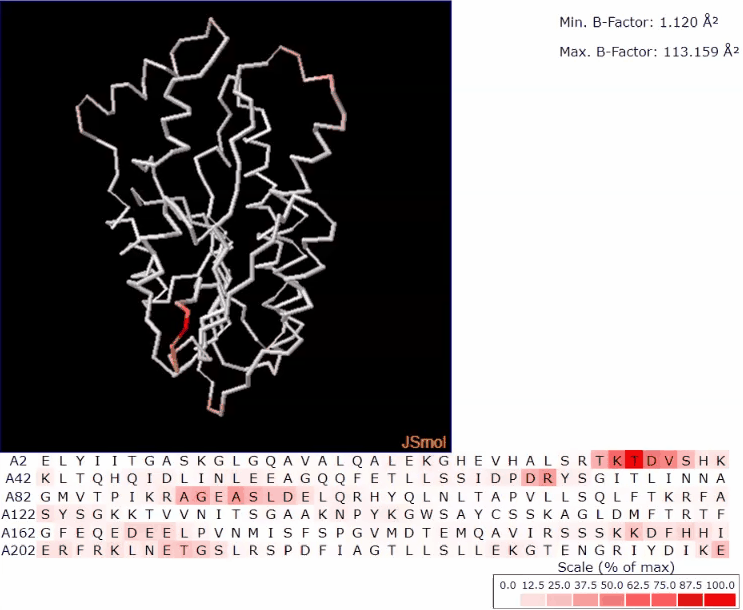

**ZRK**

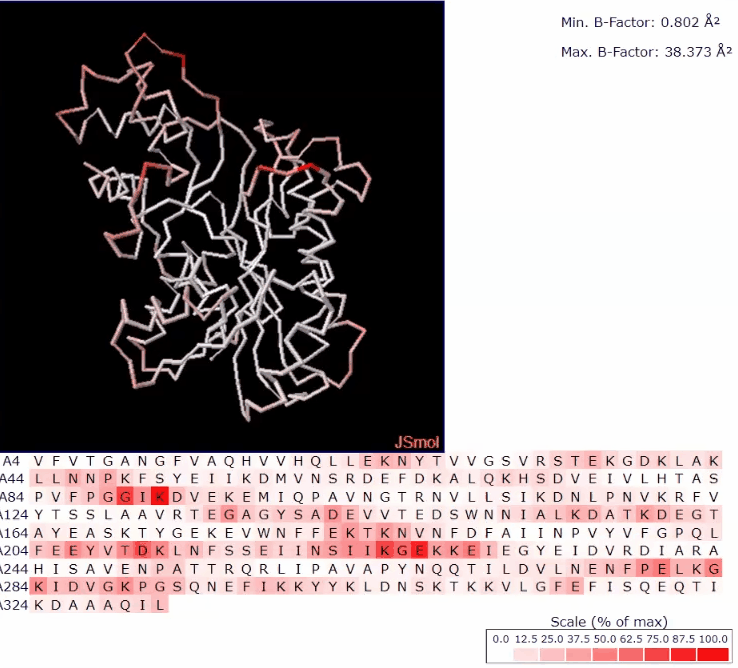

**DHK**

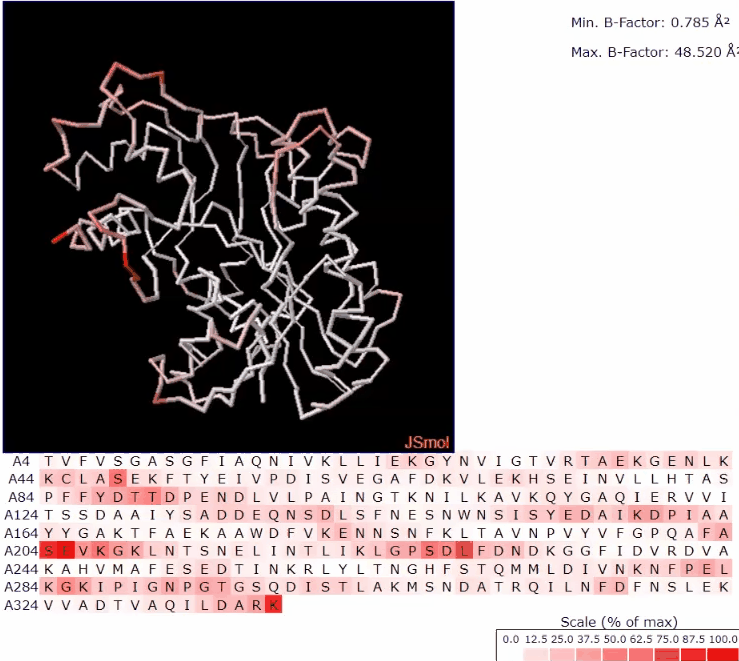

**Supplementary figure S5:** Diagram depicting the elbow method used for finding the optimal number of clusters for using the k-Means algorithm was found to be 20

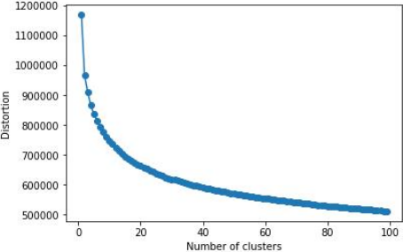

**Supplementary figure S6:** Kinetics (using Michaelis Menten fit) of various pro-pharmaceutical ketone intermediates (K-INT) using chosen keto reductases ketoreductases

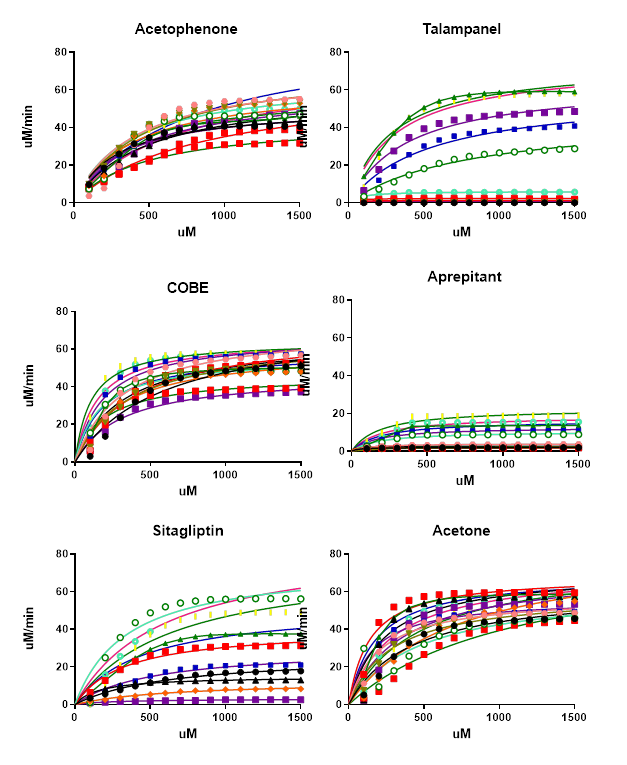

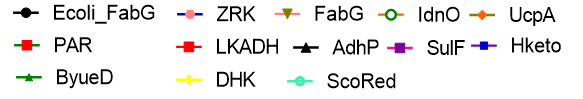

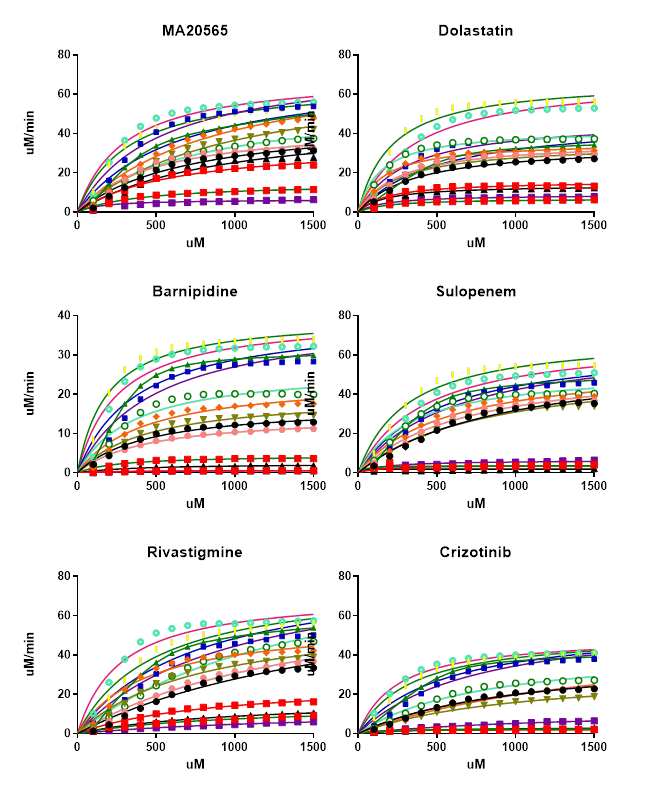

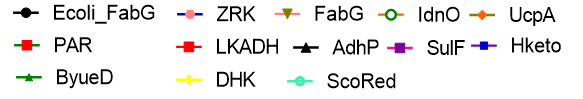

**Supplementary figure S7:** kNN (Nearest Neighbour) Test set confusion matrix

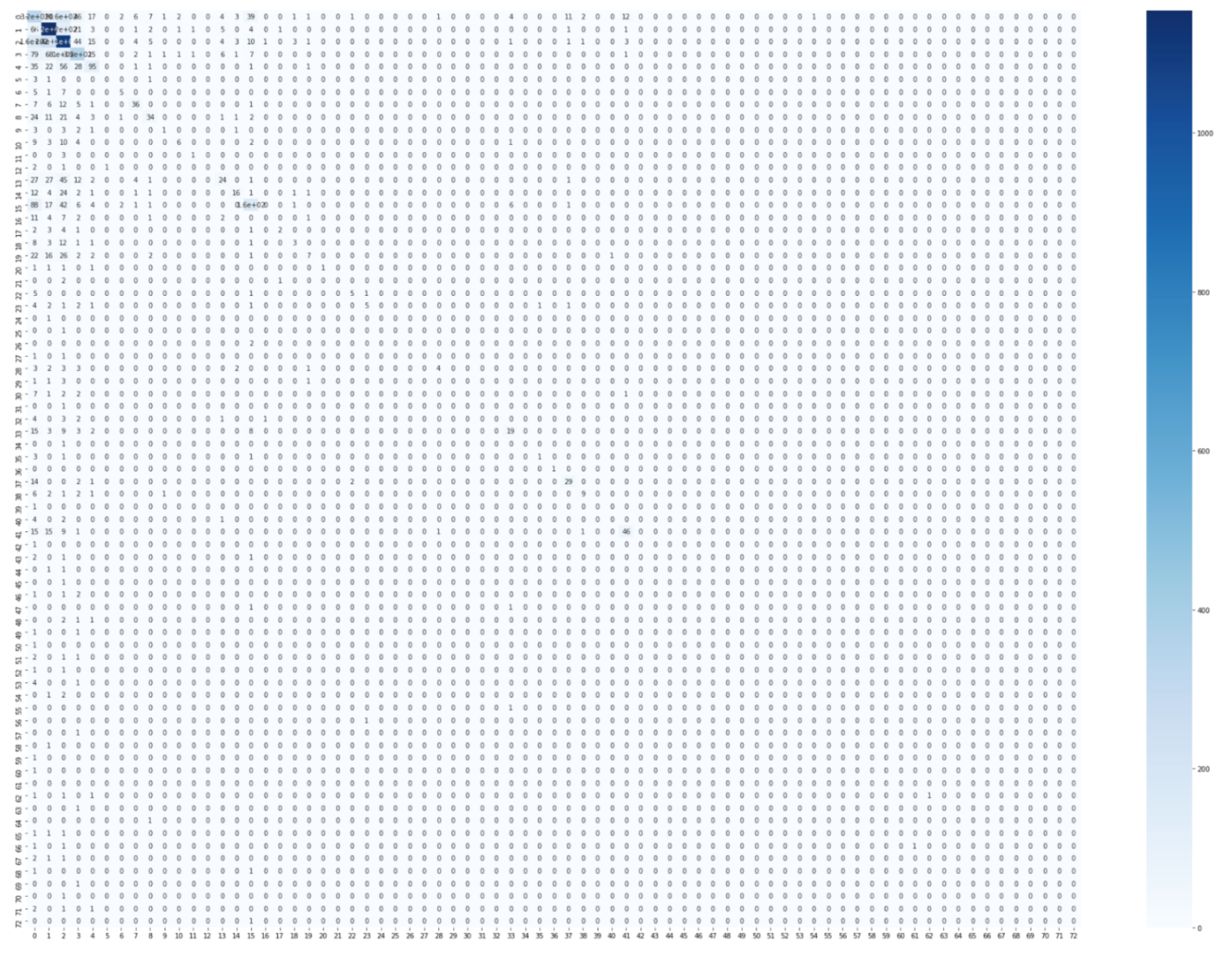

**Supplementary figure S8:** kNN (Nearest Neighbour) Training set confusion matrix
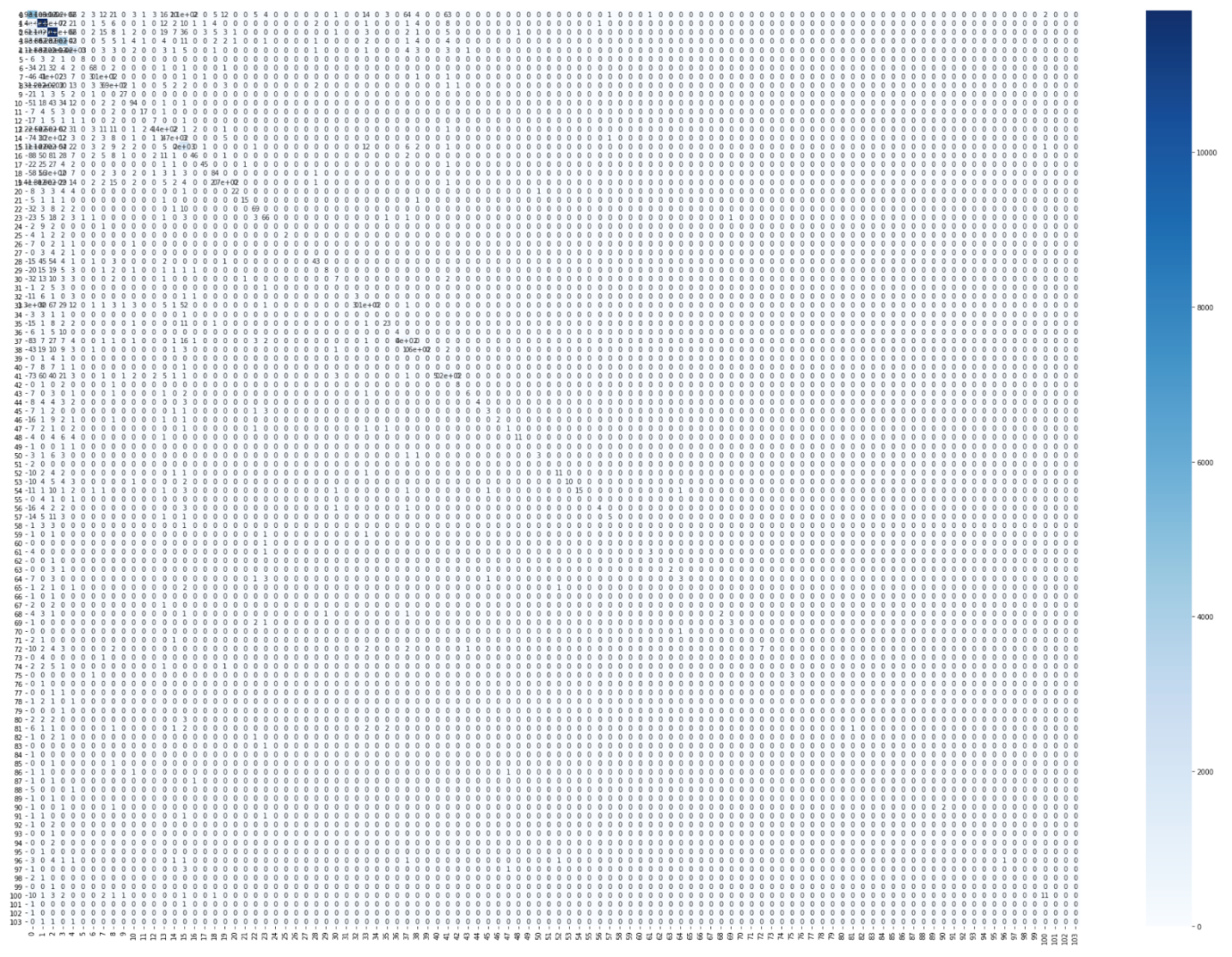

**Supplementary figure S9:** Support Vector Machines (SVM) Test set confusion matrix
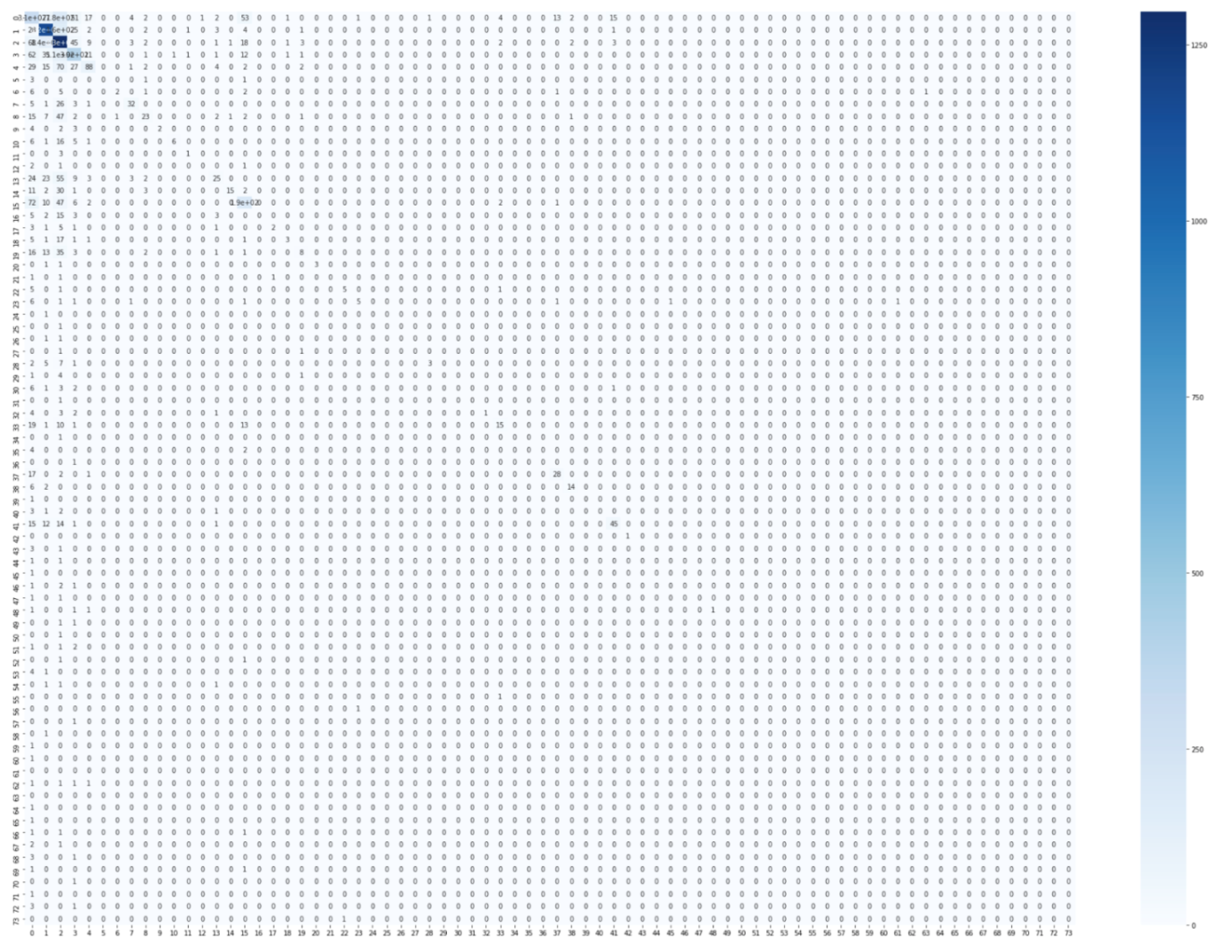

**Supplementary figure S10:** Support Vector Machines (SVM) Training set confusion matrix
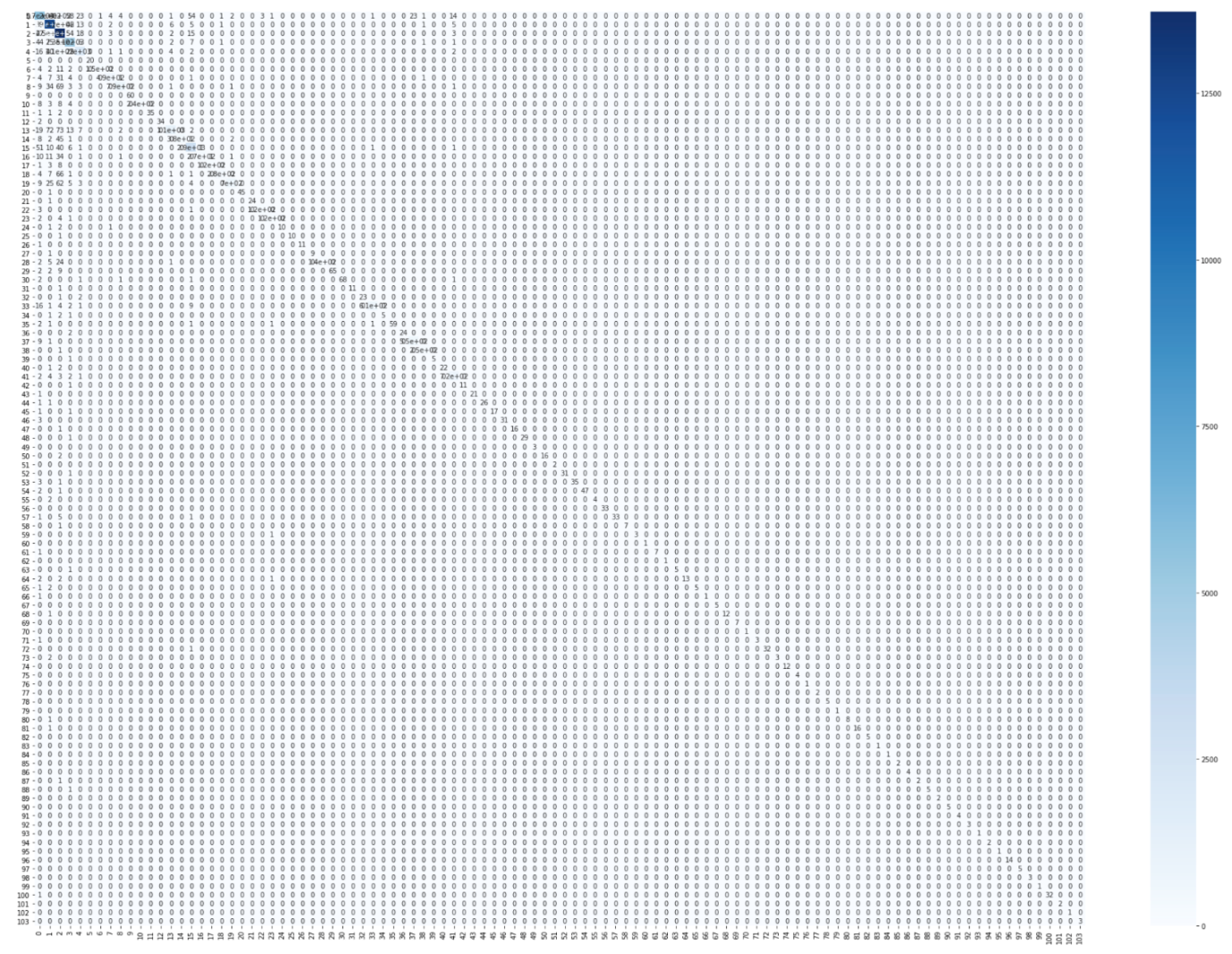

**Supplementary figure S11:** Random Forest Test set confusion matrix
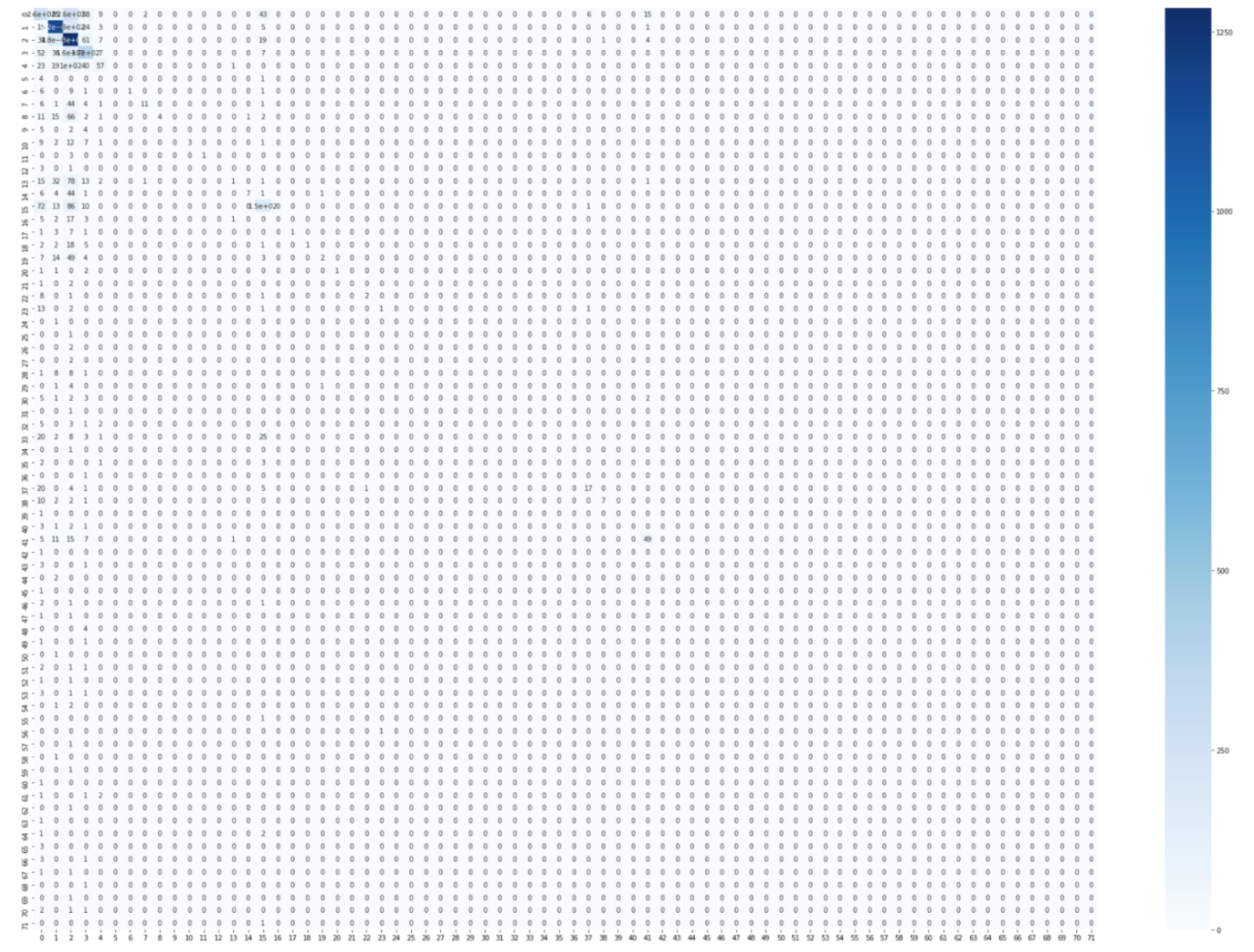

**Supplementary figure S12:** Random Forest Training set confusion matrix
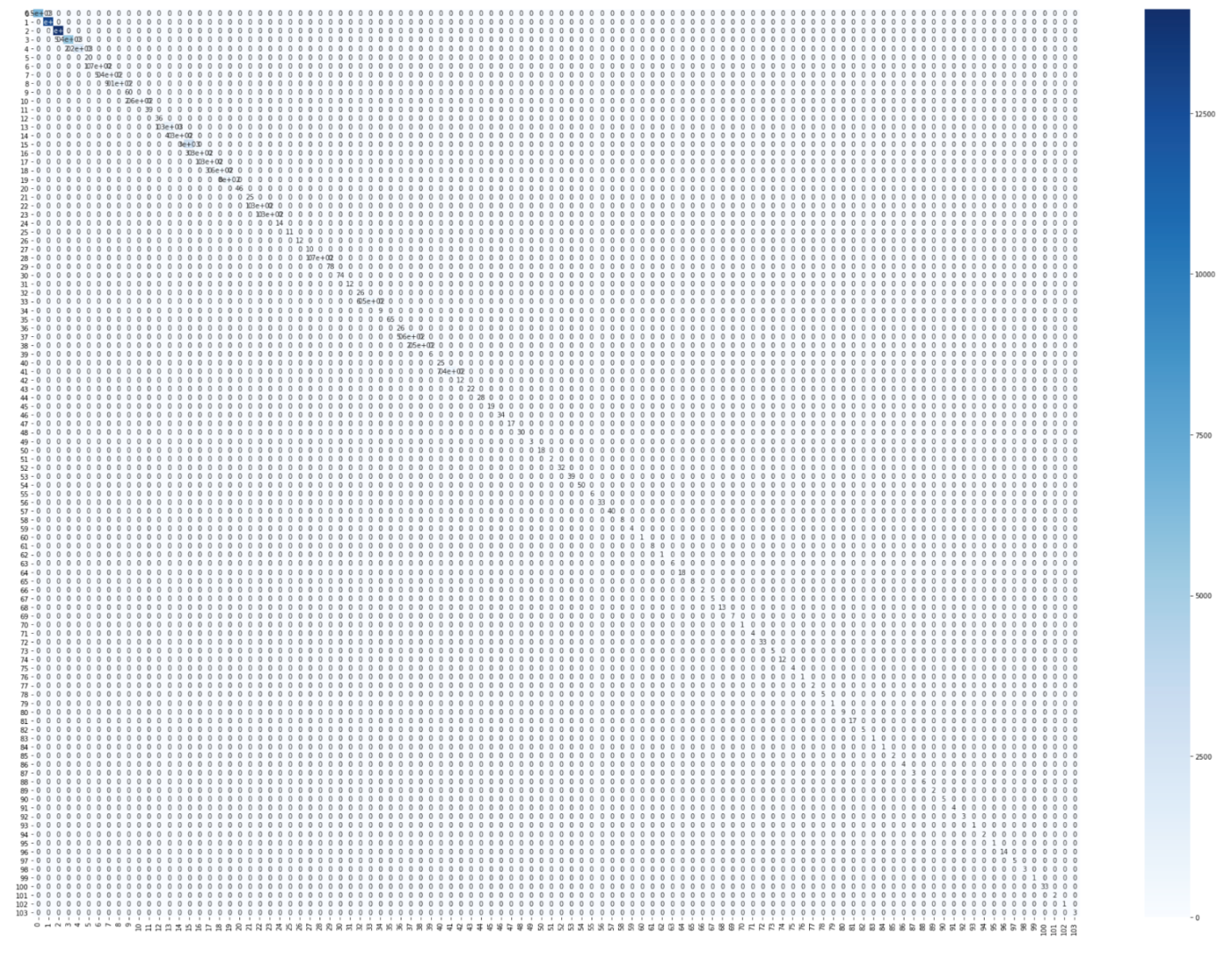

**Supplementary figure S13:** Extra Tree Test set confusion matrix

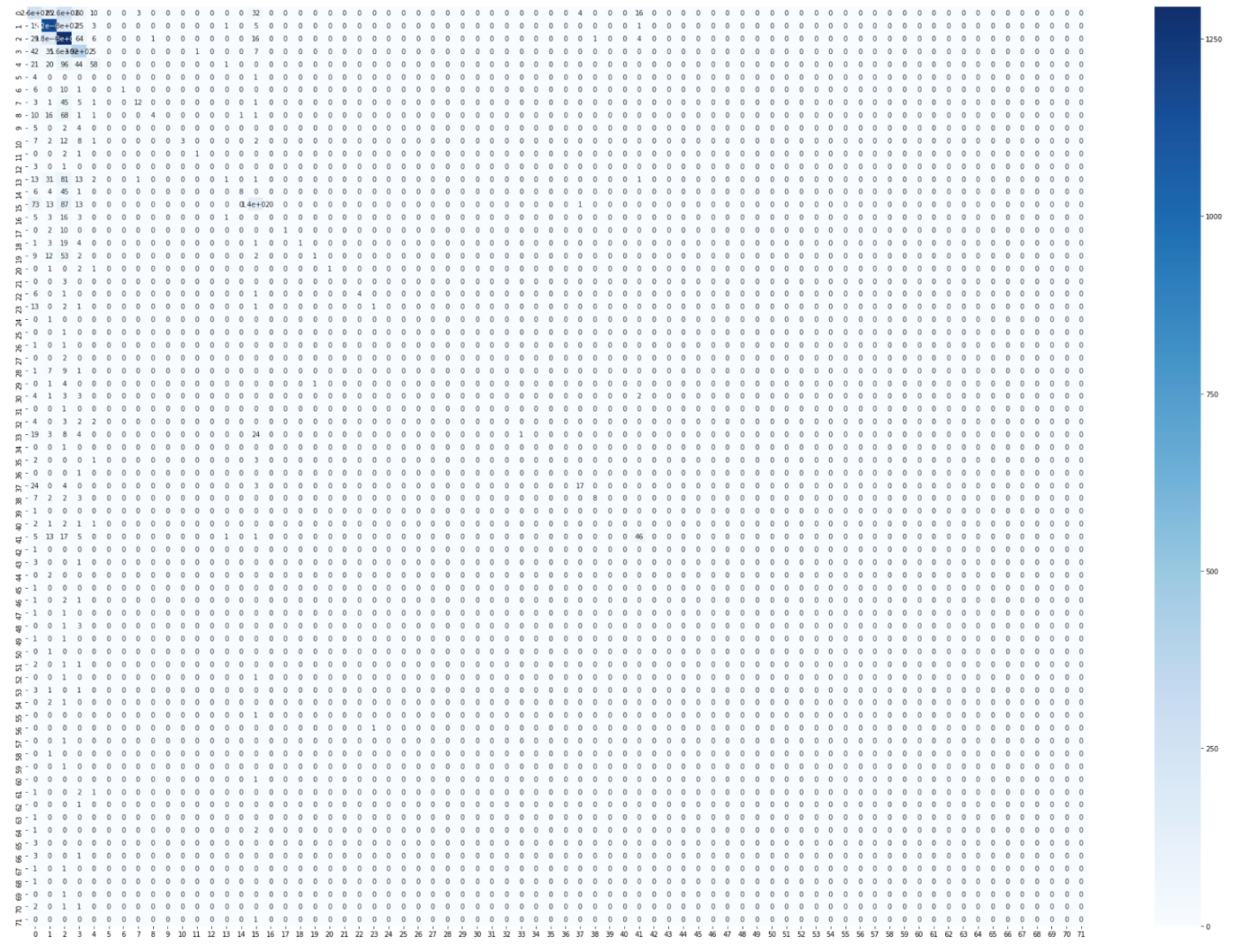

**Supplementary figure S14:** Extra Tree Test set confusion matrix

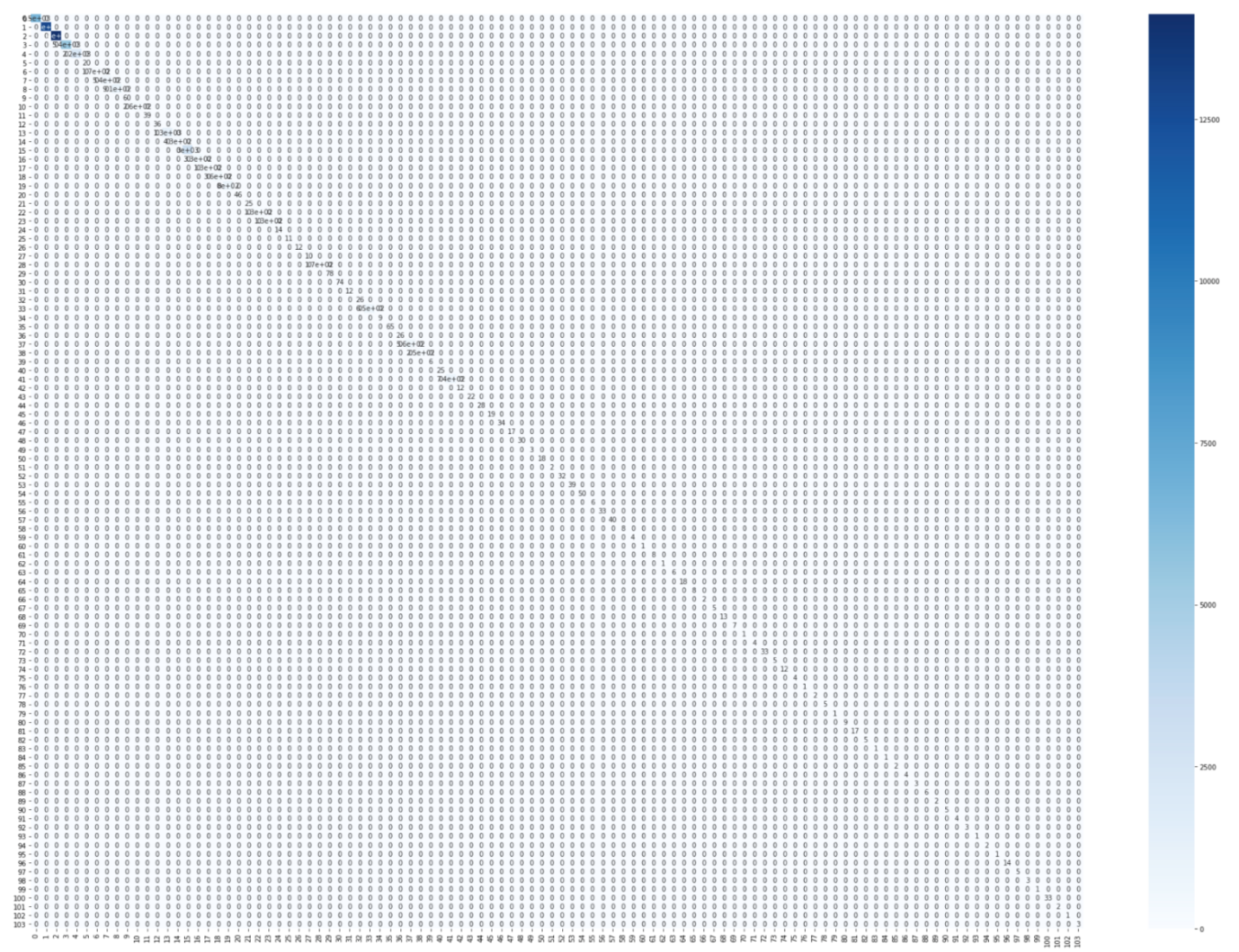
